# Genome-resolved diversity and biosynthetic potential of the coral reef microbiome

**DOI:** 10.1101/2024.08.18.608444

**Authors:** Lucas Paoli, Fabienne Wiederkehr, Hans-Joachim Ruscheweyh, Samuel Miravet-Verde, Kalia S. I. Bistolas, Teresa Sawyer, Karine Labadie, Kim-Isabelle Mayer, Aude Perdereau, Maggie M. Reddy, Clémentine Moulin, Emilie Boissin, Guillaume Bourdin, Juliette Cailliau, Guillaume Iwankow, Julie Poulain, Sarah Romac, Tara Pacific Consortium Coordinators, Serge Planes, Denis Allemand, Sylvain Agostini, Chris Bowler, Eric Douville, Didier Forcioli, Pierre E. Galand, Fabien Lombard, Pedro H. Oliveira, Jörn Piel, Olivier Thomas, Rebecca Vega Thurber, Romain Troublé, Christian R. Voolstra, Patrick Wincker, Maren Ziegler, Shinichi Sunagawa

## Abstract

Coral reefs are marine biodiversity hotspots that provide a wide range of ecosystem services. They are also reservoirs of bioactive compounds, many of which are produced by microbial symbionts associated with reef invertebrate hosts. However, for the keystone species of coral reefs—the reef-building corals themselves—we still lack a systematic assessment of their microbially encoded biosynthetic potential, and thus the molecular resources that may be at stake due to the alarming decline in reef biodiversity and cover. Here, we analysed microbial genomes reconstructed from 820 reef-building coral samples of three representative coral genera collected at 99 reefs across 32 islands during a two-year expedition throughout the Pacific Ocean (*Tara* Pacific). By contextualising our analyses with the microbiomes of other reef species, we found that genomic information was previously available for only 10% of the 4,224 microbial species overall and for less than 1% of the 645 species exclusively identified in *Tara* Pacific samples. We found reef-building coral microbiomes to be host-specific and their biosynthetic potential to rival or even surpass that found in traditional targets for natural product discovery, such as sponges and soft corals. Fire corals were not only particularly diverse in microbially encoded biosynthetic gene clusters (BGCs), but also in BGC-rich bacteria, including Acidobacteriota spp., which have been recently highlighted for their promising natural product repertoire. Together, this study unveils new candidate sources for bioactive compound discovery, prioritises targets for microbial isolation, and underscores the importance of conservation efforts by linking macro-organismal biodiversity loss to host-specific microbiomes and their biotechnological potential.

## Main text

Coral reefs are one of the most biodiverse and productive ecosystems on Earth. Despite covering less than 0.2% of the ocean floor, they are home to a third of all named marine multicellular species^1,2^. Coral reefs provide a wide range of ecosystem services to millions of people around the globe, such as food, livelihoods, and coastal protection, and they serve as a source of bioactive compounds^3,4^. However, climate change, emerging diseases, and other anthropogenic stressors have caused a decline in live coral cover by more than 50% since the 1950s^4,5^. Given the projections of further reef decline^6^, there is a pressing need to capture what is at stake under this continued biodiversity loss.

The biodiversity, productivity, and structural complexity of reef systems are fundamentally linked to the ecological functions provided by calcareous skeleton-forming (reef-building) corals, such as stony and fire corals^5^. Like other organisms, these sessile invertebrates depend on a diverse community of microorganisms (microbiome)^7^. The microbiome of corals provides its host with vital nutrients, such as carbon, nitrogen and phosphorus, as well as vitamins and essential amino acids^8–10^. Additionally, it supports its host in coping with changing environmental conditions^11^ and can protect it from infectious diseases^12,13^. Furthermore, microbes associated with reef-building corals are suggested to produce bioactive compounds to fend off pathogens, predators, and competitors^12,14^. However, such compounds have been mainly discovered in other reef invertebrates^3^, such as sponges and soft corals, and include examples of antimicrobial^15^, anti-inflammatory^16^, and antitumour^17^ agents, with some undergoing clinical trials^18,19^.

The discovery of bioactive compounds has typically relied on screening chemical extracts from either the invertebrate host, which may depend on the supply of unsustainable amounts of animal biomass^20^, or from microbial producers isolated from their hosts^17^. The latter approach is, however, generally constrained by our limited ability to cultivate microbial symbionts under laboratory conditions^21^. Furthermore, both methods are prone to the persistent challenge of rediscovering the same or similar compounds^22^. As a more recent strategy, metabolic pathways linked to biosynthetic gene clusters (BGCs) that encode the synthesis of bioactive compounds can be discovered by screening reconstructed genome sequences^23,24^. These genomes may originate from microbial culture collections^21^ as well as from uncultivated single cells or whole microbial communities (metagenomes) from, in principle, any environment or host organism^25–28^. However, for the microbiome of reef-building corals, such genomic information remains scarce^21,29^.

We thus aimed to systematically explore the genome-resolved diversity and host-specificity of reef-building coral microbiomes, assess the novelty of its BGC-encoded biosynthetic potential, compare it to the microbiomes of other hosts (such as sponges) and the surrounding environment, and determine whether corals host any BGC-rich lineages as promising targets for microbial isolation. To this end, we reconstructed >13k metagenome-assembled genomes (MAGs) from reef-building coral samples collected as part of the *Tara* Pacific expedition^30^ (Suppl. Table 1) and from publicly available coral reef metagenomic datasets (Suppl. Table 2). For almost 90% of the 4,224 microbial species in total, or >99% of those from *Tara* Pacific samples, for which we reconstructed MAGs, no genome-resolved information was previously available. Coral and sponge microbiomes were largely host-specific and we found the biosynthetic potential (per microbial species) in reef-building corals (particularly fire corals) to be as rich, or even richer than in sponges or the surrounding waters. By detecting new biosynthetically rich bacterial lineages, including some within the phylum Acidobacteriota for which the first natural products were reported only recently^31^, our work underscores exemplarily not only the value of reef-building corals from a biotechnological perspective, but also the implications of their potential loss.

### Genome-resolved resources for coral reef microbiomes

To fill the gap in the availability of microbial genome data from reef-building corals, we collected 820 metagenomes from two stony coral genera (*Porites* and *Pocillopora*) as well as fire corals (*Millepora*) around 32 islands (99 reefs) throughout the Pacific Ocean as part of the *Tara* Pacific expedition from 2016–2018^30^ (Fig. 1ab; for details on coral host lineages see^32^). Furthermore, to facilitate a comprehensive assessment of reef-building coral microbiomes for their phylogenomic novelty and biosynthetic potential, as well as to contextualise this information across different coral reef-inhabiting species (such as sponges and soft corals), we compiled a reef metagenomic dataset (Fig. 1c; Suppl. Fig. 1). This dataset included, in addition to those from the *Tara* Pacific expedition, previously available metagenomes from 412 coral samples (from 29 genera, including 22 stony and five soft coral genera, as well as from black and fire corals) and 371 sponge samples (from 32 genera) (Fig. 1d; Suppl. Table 2).

**Figure 1:**
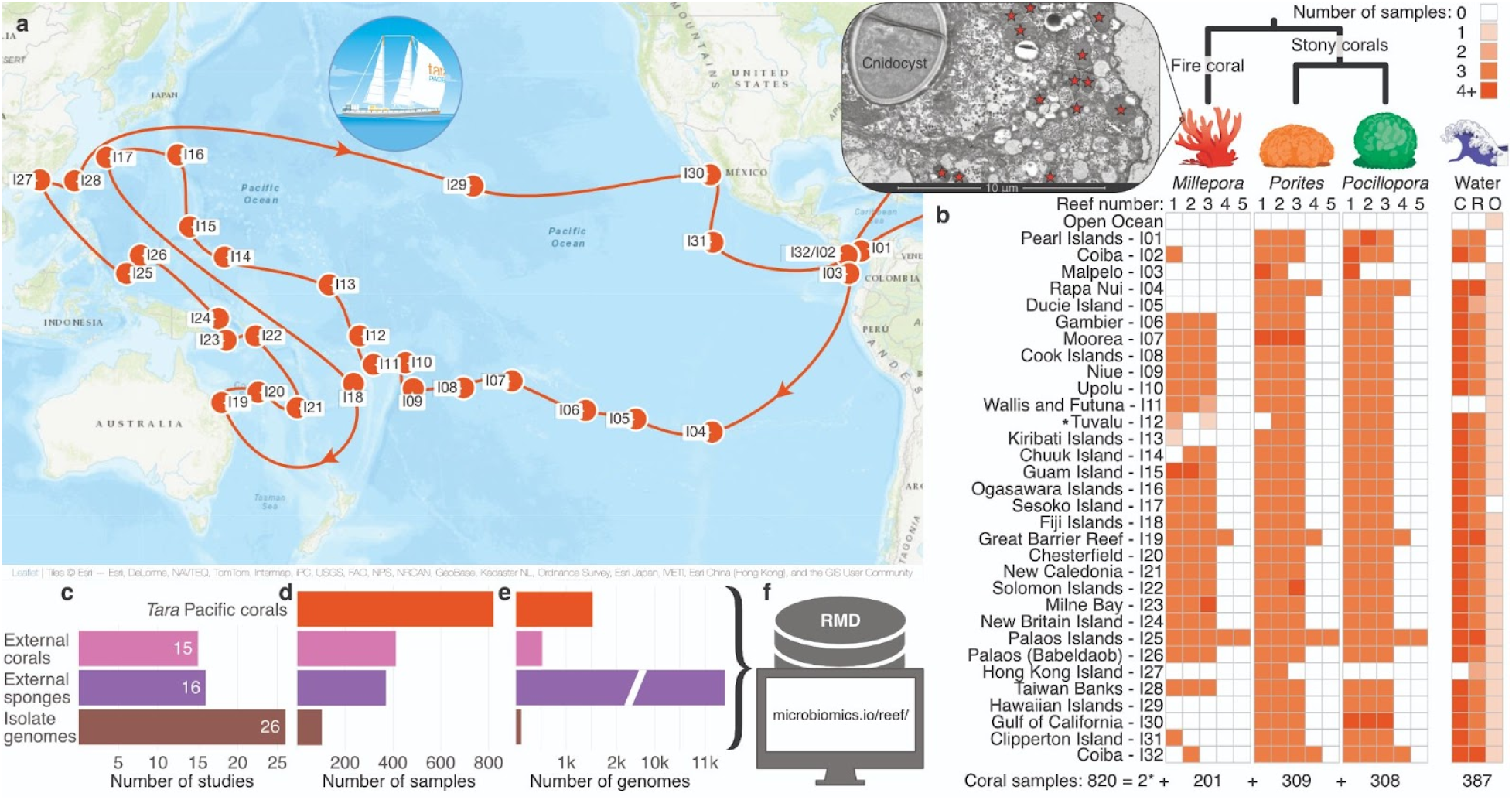
The *Tara* Pacific expedition sampled coral reefs across the Pacific Ocean. **(a)** The *Tara* Pacific expedition (2016–2018) included the sampling of corals at 99 reefs across 32 islands throughout the Pacific Ocean. **(b)** At each reef, *Millepora*, *Porites*, and *Pocillopora* colonies (*plus, exceptionally, two *Heliopora* specimens at one reef in Tuvalu) were sampled^37^, resulting in a total of 820 reef-building coral-associated metagenomes (Suppl. Table 1). In addition, the plankton microbiome was collected from the water surrounding *Pocillopora* colonies (C: coral-surrounding water), from representative water within each reef (R: reef water), as well as from oceanic water (O: open ocean water)^37^, resulting in 387 metagenomes (Suppl. Table 1). The inset shows a transmission electron microscopy image of a *Millepora* tissue sample, with bacteria-sized cells in the ectoderm indicated by stars (see Suppl. Fig. 3 and Methods for details). **(c-d)** To contextualise the data generated from the *Tara* Pacific expedition, we aggregated a total of 412 coral (from 29 genera) and 371 sponge (from 32 genera) metagenomes from 15 and 16 publicly available datasets, respectively (Suppl. Table 2). **(e)** From this metagenomic dataset, we reconstructed 13,446 metagenome-assembled genomes (MAGs), 1,524 from *Tara* Pacific metagenomes mostly from fire corals. **(c-e)** In addition, we collected 103 isolate genomes from 26 studies (Suppl. Table 4). **(f)** We compiled all genomic data to generate the Reef Microbiomics Database (RMD), which is available at https://microbiomics.io/reef/.

Applying a previously benchmarked bioinformatic workflow^28^ on this dataset, we reconstructed 13,446 coral- and sponge-associated MAGs from bacteria and archaea (Fig. 1ef, Suppl. Fig. 2). Of the 2,046 coral-associated MAGs, 1,964 were from reef-building corals and 1,524 originated from the *Tara* Pacific expedition^33^. With 57% (1,171) of all coral-associated MAGs, fire corals contributed more microbial genomes than stony (39%; 793) and soft corals (4%; 72) combined (Suppl. Table 3). Focusing on the *Tara* Pacific samples (equal sampling effort across coral hosts), we reconstructed 1,170 genomes from *Millepora* samples, 305 from *Porites* samples, and 48 from *Pocillopora* samples. Linking these genomic data to transmission electron microscopy images, we found the high number of MAGs from fire corals to correspond to a high load of extracellular microorganisms (Fig. 1g; Suppl. Fig. 3).

Overall, our efforts increased the number of available coral-associated MAGs tenfold (Suppl. Table 3). The data analysed in this study, which also includes 103 available isolate genomes (Methods; Suppl. Table 4), can be interactively explored through the Reef Microbiomics Database (RMD; https://microbiomics.io/reef/), enabling the scientific community to study the microbiomes of reef organisms.

### Novelty and host-specificity of the coral microbiome

Establishing the RMD enabled us to systematically capture the number of microbial species we reconstructed genomes for and to assess the degree to which these species lacked prior genomic information. Specifically, we annotated all genomes using the Genome Taxonomy Database (GTDB; r207) toolkit^34^ and clustered genomes using a 95% whole-genome average nucleotide-identity threshold^35^, which defined 4,224 species-level clusters (species). Close to 90% (3,774) of all species, and 99% (638) of the 645 identified in *Tara* Pacific corals, were not present in the GTDB (Fig. 2a; Suppl. Table 3). Three-quarters of the species represented in the RMD had their genomes reconstructed from sponge metagenomes (3,206 vs. 971 reconstructed from coral metagenomes) and less than 1% (36) of all microbial species were shared between sponge and coral metagenomes (Fig. 2a; Suppl. Table 3). Within corals, we reconstructed genomes from 516, 460, and 26 microbial species from stony, fire, and soft corals, respectively (Fig. 2b). Stony and fire corals shared as few as 37 microbial species, and no species were shared with soft corals. More specifically, 95% of microbial species were unique to a particular host genus (Suppl. Table 3).

**Figure 2:**
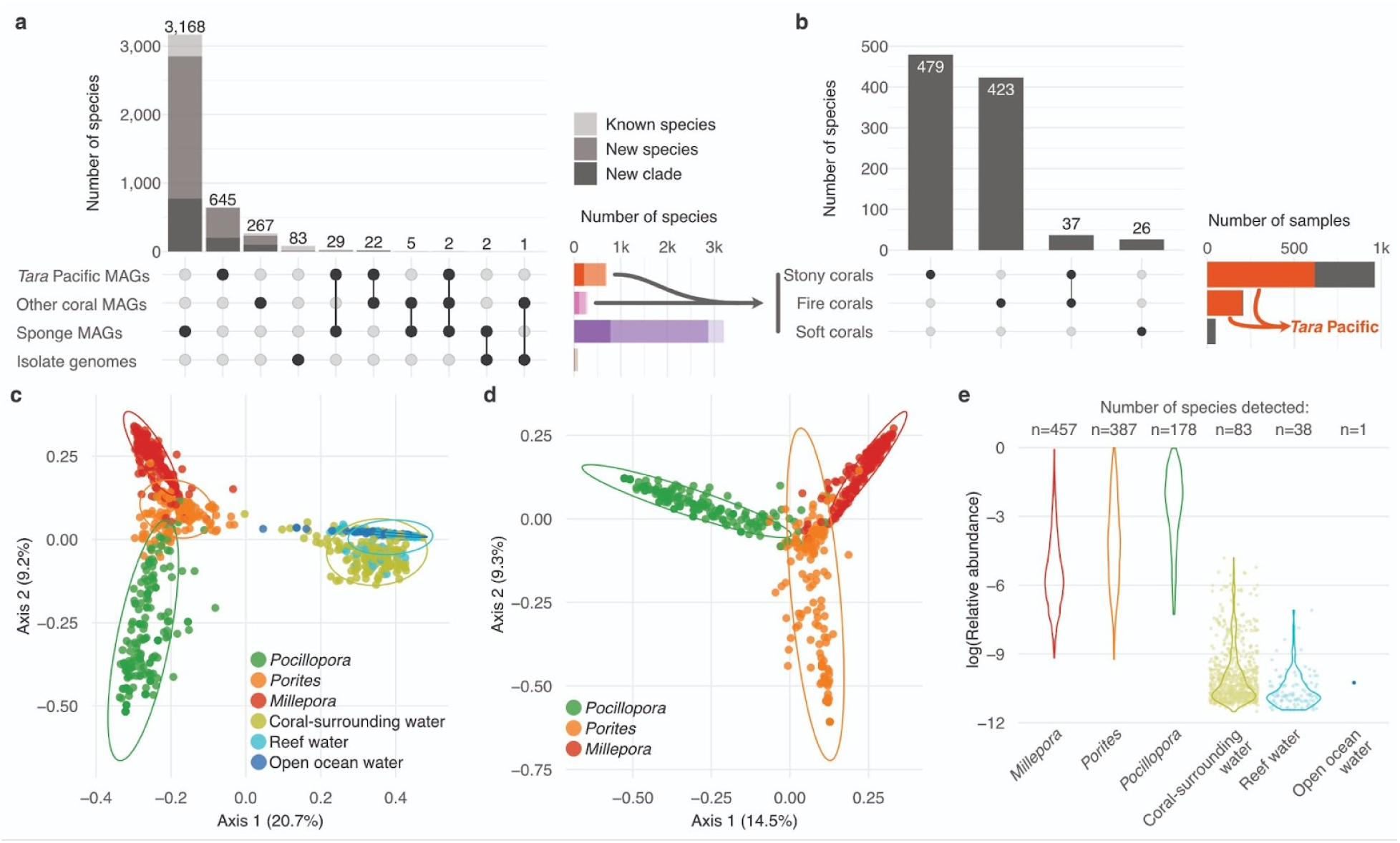
The microbiome of reef invertebrates is genomically uncharacterised and host-specific. (a) By grouping the 13,549 genomes in the RMD at the species-level (95% average nucleotide identity), we found them to represent 4,224 microbial species. Taxonomic annotation with the Genome Taxonomy Database (GTDB r207) showed that close to 90% (3,774) of all species, and 99% (638) of those reconstructed from *Tara* Pacific corals, represented new species or clades at higher taxonomic ranks. **(b)** The species displayed high specificity for the different types of corals (stony, fire, and soft corals) **(c)** A Jaccard distance-based Principal Coordinate Analysis of the microbial species we detected across the *Tara* Pacific coral and seawater metagenomes showed a clear separation between the microbiomes of corals and seawater (PERMANOVA, p-value≤0.001, R^2^=0.31). **(d)** Likewise, the coral metagenomes from the three coral genera (*Porites*, *Pocillopora*, *Millepora*) targeted during the *Tara* Pacific expedition were significantly different among themselves (PERMANOVA, p-value≤0.001, R^2^=0.18). **(e)** The number and abundance of coral-associated microbial species decreased in seawater samples as a function of distance to the coral host.

Beyond reconstructing genomes, we sought to validate this pervasive host-specificity by comparative compositional profiling of the host-associated microbial species identified here. To this end, we used a single copy marker gene-based method^36^ to determine the species-level taxonomic composition of all (1,603) host-associated metagenomes, 387 seawater metagenomes collected during the *Tara* Pacific expedition (Fig. 1b)^37^ and 84 seawater metagenomes from previous coral- and sponge-focused studies that we included in our analysis (Methods). Overall, the microbial species profiles of coral metagenomes were clearly distinct from those of seawater metagenomes (Fig. 2c and Suppl. Fig. 4) as well as among each other (Fig. 2d).

Finally, we used the *Tara* Pacific dataset to test the detectability of coral-associated microorganisms in seawater sampled at increasing distances from their host, that is, coral-surrounding water, reef water, and open ocean water. As a result, we detected as little as 20% of all microbial species from the coral metagenomes in the water samples. Furthermore, both the number and relative abundance of the detected species decreased with distance from the sampled coral colony within the reef, and they were barely detected in the open ocean (Fig. 2e).

Together, our results complement previous metagenomic efforts to reconstruct microbial genomes from diverse environments^26,38,39^, greatly expand the number and diversity of reef host-associated microbial taxa with genomic information available, and—besides enabling us to investigate their functional potential—support earlier marker-gene-based reports^40–42^ by demonstrating a high degree of host-specificity based on genome-resolved data.

### Functional and biosynthetic potential of the coral reef microbiome

To determine whether the phylogenomic novelty of the reef microbiome also correlated with an uncharted functional and biosynthetic potential, we used previously established methods^43^ to generate a reef microbial gene catalogue (RMGC) based on the protein-coding genes predicted for all genomes from the 4,224 species represented in the RMD. This resulted in 16.3 M non-redundant genes (genes), that is, a mean of 3,857 genes per species. By comparison, a recently published microbial gene catalogue from the open ocean (OMGC; derived from the Ocean Microbiomics Database (OMD))^28^, built from almost twice as many microbial species (8,304), contained substantially fewer genes per species (2,135). The higher gene richness encoded by reef host-associated microbial species was also reflected in a larger estimated genome size per species (3.6 vs 2.2 Mbp for the reef vs ocean microbiome; Suppl. Fig. 5). This difference might stem from lower streamlining pressure or vice-versa higher gene-maintenance pressure acting on microbial genomes in the nutrient-rich, complex host-associated ecological niche^44–46^. In addition, compared to the open ocean microbiome, the reef microbiome contained a higher fraction (34% vs 16%) of as-yet uncharacterised genes (Fig. 3a).

**Figure 3:**
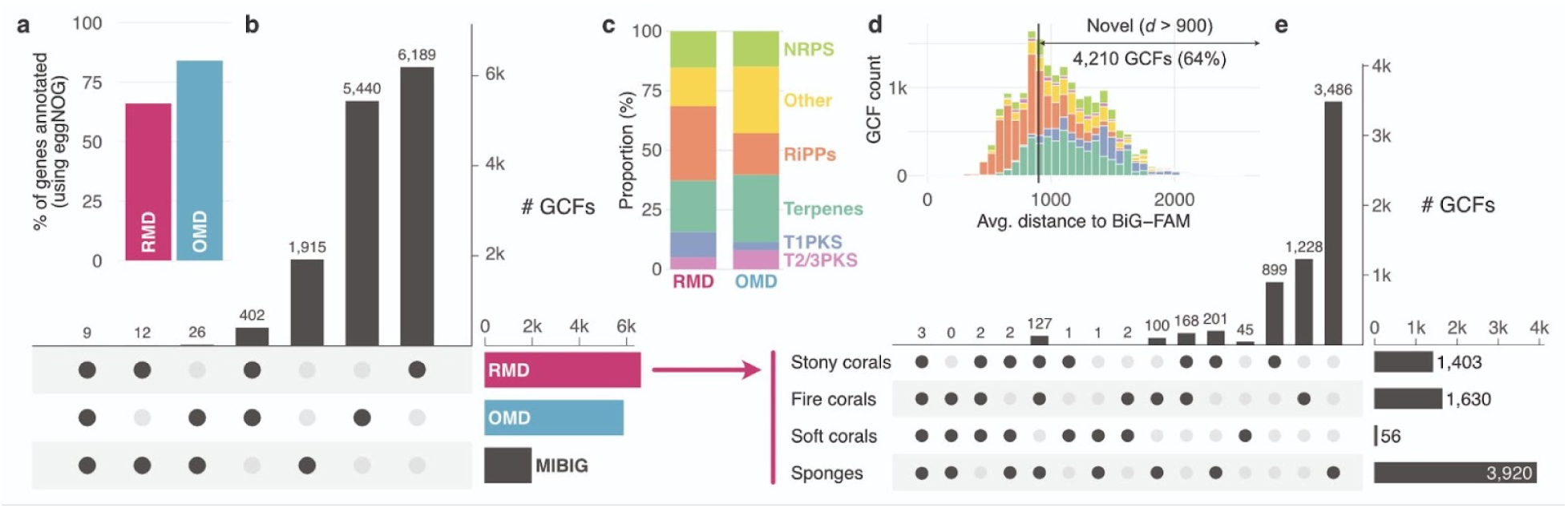
The reef microbiome harbours a high degree of functional and biosynthetic novelty. (a) The microbiomes of reef invertebrates represented in the Reef Microbiomics Database (RMD) harboured 18% fewer characterised genes than the global open ocean microbiome (gene catalogue representatives annotated using eggNOG) (RMD: 16.3M gene representatives; Ocean Microbiomics Database (OMD): 17.7M gene representatives). **(b)** We found the Gene Cluster Families (GCFs) detected in the reef microbiome to be clearly distinct from those previously detected in the open ocean^28^ and only 21 of them to overlap with biochemically characterised ones (MIBiG). **(c)** The proportion of predicted natural products for the reef and open ocean microbiomes reveals an increased representation of Ribosomally synthesised and Post-translationally modified Peptides (RiPPs) and Type I Polyketide Synthases (T1PKS) in the reef microbiome. NRPS, Non-Ribosomal Peptide Synthetases; T2/3PKS, Type II and III Polyketide Synthases. **(d)** Most of the GCFs from the reef microbiome are new compared to previously sequenced BGCs (BiG-FAM, Methods). **(e)** The GCFs detected in reef invertebrates are largely specific to the different host types (stony, fire, soft corals, and sponges).

For the microbial genome collection in the RMD, we predicted BGCs with antiSMASH (35 k BGCs from 13 k genomes) to map and contextualise the reef biosynthetic potential and to compare its magnitude and diversity with that found in open ocean microbial genomes (40 k BGCs from 35 k genomes). To ensure comparability, we grouped the antiSMASH-predicted BGCs into biosynthetic gene-cluster families (GCFs), identified pathways predicted to encode similar natural products^47^, and accounted for the inherent redundancy (the same BGC encoded in several genomes) as well as the fragmented nature of metagenomic assemblies (Methods)^48^. We found that the reef microbiome encoded a richer (6,612 GCFs or 1.57 GCF/species) biosynthetic potential than the ocean microbiome (5,877 GCFs or 0.71 GCF/species) (Fig. 3b). The two environments shared less than 10% of GCFs (411), and the proportions of the detected natural product classes also varied. For example, the reef microbiome contained a higher proportion of ribosomal natural product pathways (0.31 vs 0.174) and Type I polyketide synthases (0.107 vs 0.036), and a lower share of terpenes (0.215 vs 0.282) and other classes (0.161 vs 0.279), including aryl polyene, homoserine, beta lactones, or siderophores (Fig. 3c). These shifts may represent adaptations in the metabolism of reef invertebrate-associated microorganisms compared to free-living ones, and are in contrast to previous reports that found relatively fewer ribosomal natural products pathways in sponges compared to aquatic ecosystems (Suppl. Table 5)^26^.

Comparing the GCFs detected in coral reef microbiomes to biochemically characterised (MIBiG^49^) and predicted (BiG-FAM^50^) pathways, we found as little as 0.3% (21) overlap (Fig. 3b) and 64% to be newly identified in this study (Fig. 3d). To further support the notion that reef host-associated microorganisms represent a rich and unique source to discover new biosynthetic enzymes and natural products, we tested whether GCFs found in coral-associated microorganisms were also detected in the reef seawater; which, however, was the case for only 25% of them (which echoes the 20% overlap between corals and seawater microbiomes at the level of microbial species).

Given that reef-building corals, as opposed to sponges (followed by soft corals), have received relatively little attention as sources of marine natural products^51,52^, we compared the biosynthetic potential of the RMD across these different groups of hosts. Overall, we recovered fewer GCFs from corals than from sponges (2,817 vs 3,920 GCFs; Fig. 3e). However, once we normalised by sampling effort, the coral microbiome was richer in GCFs both per genome (2.9 vs 1.2 GCFs/species) and per microbial species (1.4 vs 0.3 GCFs/species). In addition, although the coral microbiome encoded fewer ribosomal natural product pathways (25% of all GCFs in corals compared to 36% in sponges), the coral microbiome harboured a larger share of the biotechnologically important class of nonribosomal peptide synthetases^53^ than the sponge microbiome (25% compared to 9%).

Among corals, our analyses revealed the fire coral microbiome (which has so far only been taxonomically characterised^7,54^) to be particularly BGC-rich. We detected almost twice the number of GCF per species in fire corals (4.0 GCF/species) than in soft corals (2.1 GCF/species), with stony corals (2.7 GCF/species) in-between. Although, the RMD still contains relatively few genomes from soft corals (72 MAGs) and other coral reef species, our results highlight that the microbiome of reef-building corals, and fire corals in particular, encodes an immense and yet untapped source of novel biosynthetic enzymes and natural products (Suppl. Fig. 6–8).

### BGC-rich microbial lineages in reef-building corals

Biosynthetically rich microorganisms (BGC-rich; ≥15 BGCs) are particularly valuable targets for isolating natural products. Such talented producers (or superproducers) hold tremendous biotechnological promise^28,55^ and isolating their natively produced metabolites is experimentally advantageous to subcloning and heterologously expressing single BGCs, particularly for the biotechnologically important, large multi-modular polyketide synthases and non-ribosomal peptide synthetases^56^. Previous natural product research revealed, for instance, *Candidatus* Entotheonella and *Candidatus* Poriflexus as sponge-associated BGC-rich lineages^57,58^. To identify such BGC-rich microbial lineages in our reef dataset, we screened the coral and sponge metagenomes: we mapped the taxonomic distribution of the biosynthetic potential across the RMD and, after manual curation (Methods), identified 33 BGC-rich microbial species (three times more than across the OMD^28^) from seven phyla, including Acidobacteriota, Cyanobacteria, Proteobacteria, and Myxococcota (Fig. 4a; Suppl. Table 6). However, none of these 33 BGC-rich species was represented in the GTDB, suggesting these as candidate new species (and 16 of these even candidate new genera and one a candidate new family).

**Figure 4:**
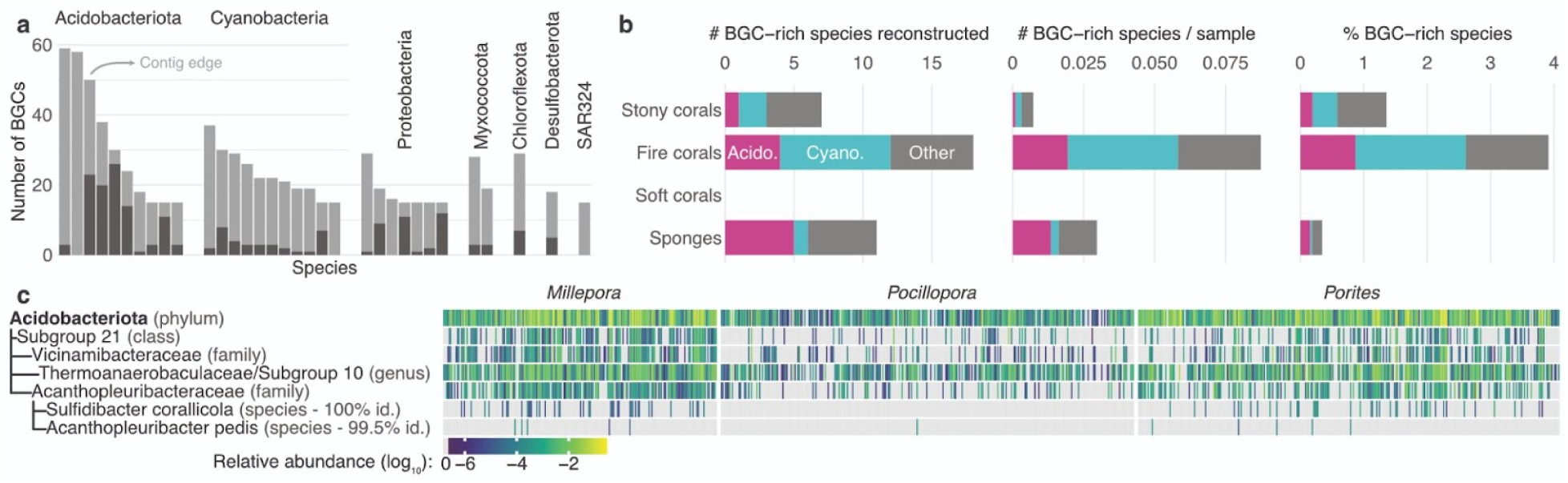
Stony and fire corals harbour BGC-rich microbial lineages. **(a)** Taxonomy and biosynthetic potential of the 33 candidate natural product superproducer species (that is, BGC-rich lineages) identified in the RMD. None of these species have representatives in the GTDB (r207). Light grey indicates BGCs located at the edge of a contig (Methods). **(b)** Fire corals host a particularly high number of BGC-rich species compared to other reef invertebrates (and we detected none in soft corals), a difference even more striking when normalising by the sampling effort. Compared to both sponges and soft corals, the microbiomes of reef building corals host a higher proportion of BGC-rich species. **(c)** We used the *Tara* Pacific 16S rRNA gene amplicon dataset (812 matching samples) to explore the abundance of Acidobacteriota and the taxonomic groups containing BGC-rich species (that is, the Subgroup 21 class, the Vicinamibactraceae and Acanthopleuribacteraceae families, and the Thermoanaerobaculaceae/Subgroup 10 genus) based on taxonomic annotations (Methods). In addition, we detected close relatives of the two isolated strains from the Acanthopleuribacteraceae family (*Acanthopleuribacter pedis* and *Sulfidibacter corallicola*), which recently led to the characterisation of phorbactazoles, 21-methoxy calyculinamide, and acidobactamides as the first acidobacterial natural products^31^.

Similar to the biosynthetic potential, we compared the distribution of BGC-rich microbial lineages across stony corals, fire corals, soft corals, and sponges. Our analysis of hundreds of metagenomes and thousands of microbial genomes showed that reef-building corals harbour a higher proportion of BGC-rich lineages (1.4% and 3.9% of all species for stony and fire corals, respectively) than soft corals (none identified) or sponges (0.3% of all species) (Fig. 4b). Fire corals in particular host many BGC-rich lineages (18, or one every 11 metagenomes) compared to sponges (11, or one every 138 metagenomes). Based on these findings, this study reveals additional potential sources in the search for novel bioactive compounds from reef invertebrate microbiomes.

The recent attention that Acidobacteriota spp. have received as candidate natural product superproducers^26,27,55^ motivated us to assess their prevalence and abundance across reef-building corals. To this end, we explored their distribution across the *Tara* Pacific 16S rRNA amplicon dataset^7^ (812 samples) that matched the *Porites*, *Pocillopora* or *Millepora* metagenomes studied here. All Acidobacteriota spp. from which we recovered BGC-rich genomes were abundant in and prevalent across the three coral genera (Methods; Fig. 4c). We also detected with a 99.5% identity match of the 16S rRNA gene, a close relative of the *Acanthopleuribacter pedis* strain, from which we recently characterised the first acidobacterial natural products (phorbactazoles, 21-methoxy calyculinamide, and acidobactamides)^31^. Phorbactazoles and 21-methoxy calyculinamide are cytotoxins with IC_50_ in the micro- and nanomolar ranges, respectively, and synthesised by complex *trans*-acyltransferase polyketide synthase (*trans*-AT PKS) pathways, which are known to produce complex and unusual compounds^59^. Furthermore, we identified two microbial genomes (reconstructed from coral metagenomes—*Millepora* and *Goniastrea*) from uncharacterised Acanthopleuribacteraceae spp. that encode unusual *trans*-AT PKS pathways (along with many other BGCs) (Suppl. Fig. 9; Suppl. Table 6).

Together, these observations suggest that reef-building corals host biosynthetically rich bacterial lineages, including Acidobacteriota spp., that emerge as new targets for isolation given their untapped yet promising potential to yield new natural products.

### Conclusions

The systematic, basin-scale sampling effort of the *Tara* Pacific expedition facilitated access to a wealth of host-specific, previously unavailable genome-resolved information for coral-associated microbial species. Comparative analyses, enabled by establishing a reef microbiome-wide database for key coral reef organisms, revealed stony and fire corals as particularly rich in novel, microbially encoded BGCs as well as BGC-rich microbial lineages. On the one hand, this enormous, largely untapped biosynthetic potential of reef-building corals, which have so far contributed only a minimal portion of the diversity of marine natural products^3,52^, represents a promising prospect for sustainable, efficient, and non-redundant bioprospecting^29^. This outlook is particularly striking considering that our work focused on only a small fraction of the globally described number of extant reef-building coral species (current estimates: >750)^60^ and their associated microbial diversity^7^, the vast majority of which is yet genomically undescribed. On the other hand, our findings highlight what is at stake under the continued, anthropogenic pressure-driven decline of coral reefs and the unique microbial diversity they support.

## Methods

### Metagenomic data generation from *Tara* Pacific samples

The *Tara* Pacific expedition collected coral and seawater samples from 99 reefs across 32 islands throughout the Pacific Ocean^30^. The detailed sampling protocols are presented in Lombard et al.^37^. Based on morphology, we targeted three reef-building corals: the stony corals *Pocillopora meandrina* and *Porites lobata*, and the fire coral *Millepora* cf. *platyphylla*. However, because corals within the same genus can be difficult to differentiate by eye, results were aggregated at the genus level (*Pocillopora*, *Porites*, and *Millepora*). Population genomic analysis of the sampled morphotypes revealed the presence of five, three, and one putative species for *P. meandrina*, *P. lobata*, and *M.* cf. *platyphylla*, respectively^32^. In addition, two samples of *Heliopora* were collected at one site.

Samples from three colonies per species were collected for metagenomic analysis at each site using hammer and chisel. The fragments were stored underwater in individual Ziploc bags and preserved in tubes with DNA/RNA shield (Zymo Research, Irvine, CA, USA) at −20 °C once on board the schooner *Tara*. Additionally, plankton was collected on 0.2–3 µm filters by filtering 100 L of open ocean water with a pump offshore, 100 L of reef water with a pumping system and tubing within the reefs, and 50 L of coral-surrounding water with a pumping system and diver-held tubing from two *Pocillopora* colonies per reef. All filters were preserved in cryovials in liquid nitrogen. The sample provenance and environmental context are available on Zenodo^61^.

DNA was extracted with commercial kits after mechanical cell disruption and metagenomic sequencing was performed on a NovaSeq6000 or HiSeq4000 Illumina sequencer (Illumina, San Diego, CA, USA) in order to produce 100 M of ∼150 bp paired-end reads per sample. The detailed protocols are provided in Belser et al.^62^.

### Transmission electron microscopy images of coral tissue samples

The *Tara* Pacific expedition also enabled the microscopical inspection of coral tissue. For this, 0.1 g of coral was separated using bone cutters and immediately preserved on board in EM-grade glutaraldehyde. In the lab, the coral fragments were decalcified in 500 μL of 10% EDTA, pH 7. Once decalcified, the samples were rinsed with 0.1 M sodium cacodylate buffer and embedded in agarose for post-fixation staining with 1.5% potassium ferrocyanide and 2% osmium tetroxide in water. Osmium tetroxide staining was followed by uranyl acetate and lead nitrate staining^63^. Next, the samples were dehydrated in serial acetone baths (10, 30, 50, 70, 90, 95, 100%) for 10–15 min each before being infiltrated with araldite resin and sectioned using a RMC ultramicrotome. Samples were imaged on a FEI Helios Nanolab 650 microscope in scanning transmission electron microscopy mode at the Oregon State University Electron Microscopy Facility.

### Inclusion of publicly available sequence data from other corals and sponge microbiomes

We complemented the newly generated *Tara* Pacific metagenomic dataset with publicly available ones. Specifically, we searched the European Nucleotide Archive (ENA) biosamples database (December 2022) to identify metagenomic datasets from coral or sponge samples. Upon literature review, we included metagenomic data from 15 coral^64–78^ and 16 sponge^27,79–93^ focused studies (Suppl. Table 2). In total, these datasets included 412 coral (from 29 genera, including 22 stony and five soft coral genera, as well as black and fire corals) and 371 sponge samples of global distribution (Suppl. Fig. 1). An additional 103 microbial genomes from cultivated coral-associated strains, reported across 26 studies^21,51,94–113^, were included (Suppl. Table 4).

### Metagenomic data assembly and binning

Metagenomic data processing was performed as described in Paoli et al.^28^. Briefly, sequencing reads from all metagenomes were quality filtered using BBMap (v.38.79) by removing sequencing adapters from the reads, removing reads that mapped to quality control sequences (PhiX genome), and discarding low quality reads using the parameters trimq = 14, maq = 20, maxns = 1, and minlength = 45. Downstream analyses were performed using quality-controlled reads or, if specified, merged quality-controlled reads (bbmerge.sh minoverlap = 16). For host-associated metagenomes (that is, excluding seawater samples), quality-controlled reads from *Tara* Pacific metagenomes were normalised (bbnorm.sh target = 40, mindepth = 0) and all metagenomes were assembled individually with metaSPAdes (v.3.14.1 or v.3.15 if required)^114^. The resulting scaffolded contigs (hereafter scaffolds) were filtered by length (≥1 kbp).

The 1,603 metagenomic samples were grouped into several sets and, for each sample set, the quality-controlled metagenomic reads from all samples were individually mapped against the scaffolds of each sample. Publicly available metagenomes were processed by study, while the *Tara* Pacific coral metagenomes were processed by host (creating several sets of approximately 100 samples within each host genus). Reads were mapped with BWA (v.0.7.17-r1188)^115^, allowing the reads to map at secondary sites (with the -a flag). Alignments were filtered to be at least 45 bases in length, with an identity of ≥97% and covering ≥80% of the read sequence. The resulting BAM files were processed using the jgi_summarize_bam_contig_depths script of MetaBAT 2 (v.2.12.1)^116^ to provide within- and between-sample coverages for each scaffold. The scaffolds were finally binned by running MetaBAT 2 on all samples individually with parameters --minContig 2000 and --maxEdges 500 for increased sensitivity.

### Quality evaluation of metagenomic bins and additional genomes

The quality of each metagenomic bin and external genome was evaluated using both the ‘lineage workflow’ of CheckM (v.1.1.3)^117^ and Anvi’o (v.7.1)^118^. Metagenomic bins and external genomes were retained for downstream analyses if either CheckM or Anvi’o reported a completeness/completion (cpl) of ≥50% and a contamination/redundancy (ctn) of ≤10%. Completeness and contamination were averaged across both tools and the selected genomes were further attributed a quality score (Q) as follows: Q = cpl − 5 × ctn. Based on the quality score, the bins were classified as MAGs of different quality levels according to community standards^119^ as follows: high quality: cpl ≥ 90% and ctn ≤ 5%; good quality: cpl ≥ 70% and ctn ≤ 10%; medium quality: cpl ≥ 50% and ctn ≤ 10%; fair quality: cpl ≤ 50% or ctn ≥ 10% (edge case due to the average between the CheckM and Anvi’o).

### Species-level clustering of the genome collection

The genomes were subsequently grouped into species-level clusters on the basis of whole-genome average nucleotide identity (ANI) using dRep (v.3.0.0)^120^ with a 95% ANI threshold^35,121^ (-comp 0 -con 1000 -sa 0.95 -nc 0.2). Whenever mentioned, a representative genome was selected based on the maximum quality score defined by dRep (Q′ = cpl − 5 × ctn + ctn × (strain heterogeneity)/100 + 0.5 × log[N50]) for each species-level cluster.

### Taxonomic and functional genome annotation

Prokaryotic genomes were taxonomically annotated using GTDB-Tk (v.2.1.0)^34^ with the default parameters and the fulltree option using the GTDB r207 release^122^. The taxonomic annotation of a species is defined as the one of its representative genome.

Each genome was functionally annotated by first predicting complete genes using Prodigal (v.2.6.3)^123^, Barrnap (v.0.9) and Aragorn (v.1.2.41)^124^ with the taxonomy parameter specified based on GTDB annotations, where appropriate. The predicted proteins were used to identify universal single-copy marker genes with fetchMGs (v.1.2)^125^. The gene sequences were additionally used as input to identify BGCs in the genomes using antiSMASH (v.6.1.1)^23^ and a minimum contig length of 5 kbp.

### Species-level profiling with mOTUs

Genomes were added to the database (v.3.1) of the metagenomic profiling tool mOTUs^125^ to generate an extended mOTUs reference database. Only genomes with at least six out of the ten universal single copy marker genes were kept. The resulting 10,511 genomes grouped into 3,673 mOTUs species-level clusters, 3,588 (98%) of which represented new clusters in the extended database. The difference in the number of species-level clusters reported by mOTUs (3,673) and dRep (4,224) can be explained by a more stringent filtering of the genomes (10,511 instead of 13,549) and a different clustering strategy, yet, these two independent methods yielded very similar clustering patterns with a V-measure of 0.99, a homogeneity of 0.99, and a completeness of 0.98. Profiling of the 2,074 metagenomes was performed using the default parameters of mOTUs (v.3.1).

To assess the distribution (detection, abundance) of microbial species across host and seawater samples, the resulting profiles were filtered to retain the 3,673 mOTUs for which genomes were available. We subsequently removed mOTUs that were detected in less than 10 samples and samples that had a total scaled abundance of three or less. The resulting matrix was used to compute Jaccard distances between samples. These were visualised by a Principal Coordinate Analysis (PCoA) (package ape^126^ in R) and the statistical significance was tested using PERMANOVA. In case of unbalanced groups (for instance, 337 sponge metagenomes vs 47 sponge-associated seawater metagenomes), a random subsample (here 50) of the larger group was selected to perform the PERMANOVA and the statistics were derived from 100 repetitions.

### Functional profiling

Similar to the methods described previously^28,43,127^, we clustered the >37 million protein-coding genes at 95% identity and 90% coverage of the shorter gene using CD-HIT (v.4.8.1)^128^ into 16.3 million gene clusters. The longest sequence was selected as the representative gene of each gene cluster. The representative genes were annotated by assigning them to orthologous groups with emapper (v.2.1.7)^129^ based on eggNOG (v.5.0.2)^130^, and by performing queries against the KEGG database (release 2022-04)^131^. This last step was performed by aligning the corresponding protein sequences using DIAMOND (v.2.0.15.153)^132^ with a query and subject sequence coverage of ≥70%. The results were further filtered on the basis of the bitscore being ≥50% of the maximum expected bitscore (query sequence aligned against itself) in accordance with the thresholds implemented in the NCBI Prokaryotic Genome Annotation Pipeline^133^.

The 1,603 host-associated and 471 seawater metagenomes were then mapped to the cluster representatives with BWA (v.0.7.17-r1188) (-a) and the resulting BAM files were filtered to retain only alignments with a percentage identity of ≥95%, an alignment length of ≥80% of the query length and ≥45 bases. Gene abundance was calculated by first counting inserts from best unique alignments and then, for ambiguously mapped inserts, adding fractional counts to the respective target genes in proportion to their unique insert abundances. These abundance profiles were subsequently length-normalised based on the length of the gene representatives and sequencing depth-normalised using the total mOTUs counts from the corresponding sample.

### Clustering of BGCs into GCFs

All BGCs predicted across all genomes, along with those from the OMD^28^ and MIBiG^49^ (for contextualisation), were grouped into Biosynthetic Gene Cluster Families (GCFs) using clust-o-matic with a threshold of 0.5^134^ based on the comparison of biosynthetic genes. This sequence identity-based clustering was previously shown to perform similarly to BiG-SLiCE^47^, yet to provide an alternative that is less sensitive to BGC fragmentation. Each GCF was attributed a natural product class on the basis of antiSMASH annotations of the individual BGCs, as defined in BiG-SCAPE^135^. The fraction of new GCFs was estimated based on the distances to databases of computationally predicted (the RefSeq database within BiG-FAM)^50^ and experimentally validated (MIBiG 2.0)^49^ BGCs.

### *Acanthopleuribacter pedis* and *Sulfidibacter corallicola* genomes

The genomes of *A. pedis* and *S. corallicola* were downloaded from the NCBI genome database using the accessions GCF_017377855.1 and GCF_017498545.1, respectively.

### *Tara* Pacific 16S rRNA dataset

To further explore the distribution of selected microbial lineages across the coral hosts sampled by *Tara* Pacific, we used the *Tara* Pacific 16S rRNA gene (16S) dataset (v1.1)^7^ available on Zenodo (https://zenodo.org/record/4451892)^136^ and selected the 812 samples that matched the 820 coral metagenomes. The species for which MAGs were reconstructed in this study were matched to the 16S-based OTUs by (1) recovering, whenever available, the genomic 16S sequences directly from the genomes or (2) identifying the 16S sequences from related representatives in the GTDB and matching them to the 16S-based OTUs. These sequences enabled us to match whole-genome and 16S-based taxonomies to identify the corresponding 16S-based OTUs belonging to candidate BGC-rich lineages.

### Statistics and reproducibility

Statistical analyses were performed in R (v.4.2.2-4.3.1). Wherever appropriate, correction for multiple testing was performed using false-discovery rate correction. Wherever appropriate and if not specified otherwise, statistical tests performed were two-sided. UpSet plots were generated using the R packages UpSetR (v.1.4.0)^137^ and ComplexUpset (v.1.3.3)^138,139^. Boxplots were generated using ggplot2 (v.3.4.2) and defined as follows: the bottom and top hinges correspond to the first and third quartiles (the 25th and 75th percentiles). respectively, the top and bottom whiskers extend from the hinge to the largest or smallest value no further than 1.5 × IQR from the hinge, respectively (where the IQR is the interquartile range, or distance between the first and third quartiles). Outliers (data points beyond the end of the whiskers) are plotted individually. Wherever possible, individual data points are displayed, except, for instance, for coral samples in Figure 2e, owing to the large number of points and space constraints.

## Data availability

The metagenomic data generated in this study was submitted to the ENA at the EMBL European Bioinformatics Institute under the *Tara* Pacific Umbrella project PRJEB47249. Sample provenance and environmental context are available on Zenodo (https://zenodo.org/doi/10.5281/zenodo.4068292)^61^. The publicly available metagenomic data used in this study were downloaded from the ENA and a summary of their accession numbers is provided in Supplementary Table 1. The MIBiG and BiG-FAM databases can be accessed at https://mibig.secondarymetabolites.org/ and https://bigfam.bioinformatics.nl/, respectively. Other supporting data were deposited on Zenodo (https://zenodo.org/doi/10.5281/zenodo.10182966)^140^ and the RMD can be interactively accessed online (https://microbiomics.io/reef/). Additional material generated in this study is available on request.

## Code availability

The code used for the analyses performed in this study is accessible at GitHub (https://github.com/SushiLab/reef-microbiomics-paper/) and archived on Zenodo (https://zenodo.org/doi/10.5281/zenodo.10201847).

## Acknowledgments

Special thanks to the *Tara* Ocean Foundation, the R/V *Tara* crew, and the *Tara* Pacific Expedition Participants (https://doi.org/10.5281/zenodo.3777760). We are keen to thank the commitment of the following institutions for their financial and scientific support that made this unique 3-year *Tara* Pacific Expedition possible: CNRS, PSL, CSM, EPHE, Genoscope, CEA, Inserm, Université Côte d’Azur, ANR, agnès b., the Veolia Foundation, the Prince Albert II de Monaco Foundation, Région Bretagne, Lorient Agglomération, L’Oréal, Biotherm, Billerudkorsnas, AmerisourceBergen Company, Oceans by Disney, France Collectivités, Fonds Français pour l’Environnement Mondial (FFEM), UNESCO-IOC, Etienne Bourgois, and the *Tara* Ocean Foundation teams. *Tara* Pacific would not exist without the continuous support of the participating institutes. We would like to express our deepest gratitude towards all the countries and local authorities that have enabled and supported the sampling conducted as part of the *Tara* Pacific expedition (Suppl. Information). The sampling permits associated with the data generated as part of this study are available as Supplementary Table 7. Although the sequence information associated with these samples has been archived on public repositories according to best practices in the field, it should be noted that any downstream commercial use should be done in accordance with international law (Suppl. Table 7). This study was supported in part by FRANCE GENOMIQUE (ANR-10-INBS-09). S.S. acknowledges the support of the Swiss National Science Foundation (project grants: 205321_184955, 205320_215395 and the NCCR Microbiomes: 51NF40_180575) and support from the ETH IT services for computations on the ETH Euler cluster. S.M.V was supported by the Human Frontiers Science Program (HFSP) Long-term Fellowship (LT0050/2023-L1). The authors also particularly thank Serge Planes, Denis Allemand, and the *Tara* Pacific consortium. This is publication number #XX of the *Tara* Pacific Consortium.

## *Tara* Pacific Consortium Coordinators

Emilie Boissin^8^, Serge Planes^8,10^, Denis Allemand^12,13^, Sylvain Agostini^14^, Chris Bowler^15^, Eric Douville^16^, Didier Forcioli^17^, Pierre E. Galand^18^, Fabien Lombard^10,19^, Olivier Thomas^6^, Rebecca Vega Thurber^2^, Romain Troublé^7,10^, Christian R. Voolstra^5^, Patrick Wincker^4,10^, Maren Ziegler^21^, Shinichi Sunagawa^1^, Bernard Banaigs^8^, Colomban de Vargas^11^, J. Michel Flores^23^, Paola Furla^12^, Eric Gilson^24^, Stéphane Pesant^25^, Stephanie Reynaud^12^, Didier Zoccola^12^

^23^Weizmann Institute of Science, Department of Earth and Planetary Sciences, Rehovot, Israel

^24^Université Côte d’Azur, CNRS, INSERM, Institute for Research on Cancer and Aging, Nice (IRCAN), Nice, France

^25^European Molecular Biology Laboratory, European Bioinformatics Institute, Wellcome Genome Campus, Hinxton, United Kingdom

## Supplementary Tables

Supplementary Table 1 - *Tara* Pacific metagenomes

Supplementary Table 2 - Publicly available metagenomes from coral and sponge studies

Supplementary Table 3 - Reef Microbiomics Database (RMD) summary Supplementary Table 4 - Isolate genomes included in this study Supplementary Table 5 - Biosynthetic potential of the RMD genomes Supplementary Table 6 - Candidate BGC-rich lineages

Supplementary Table 7 - Summary of the sampling permits

---

**Supplementary Figure 1:**
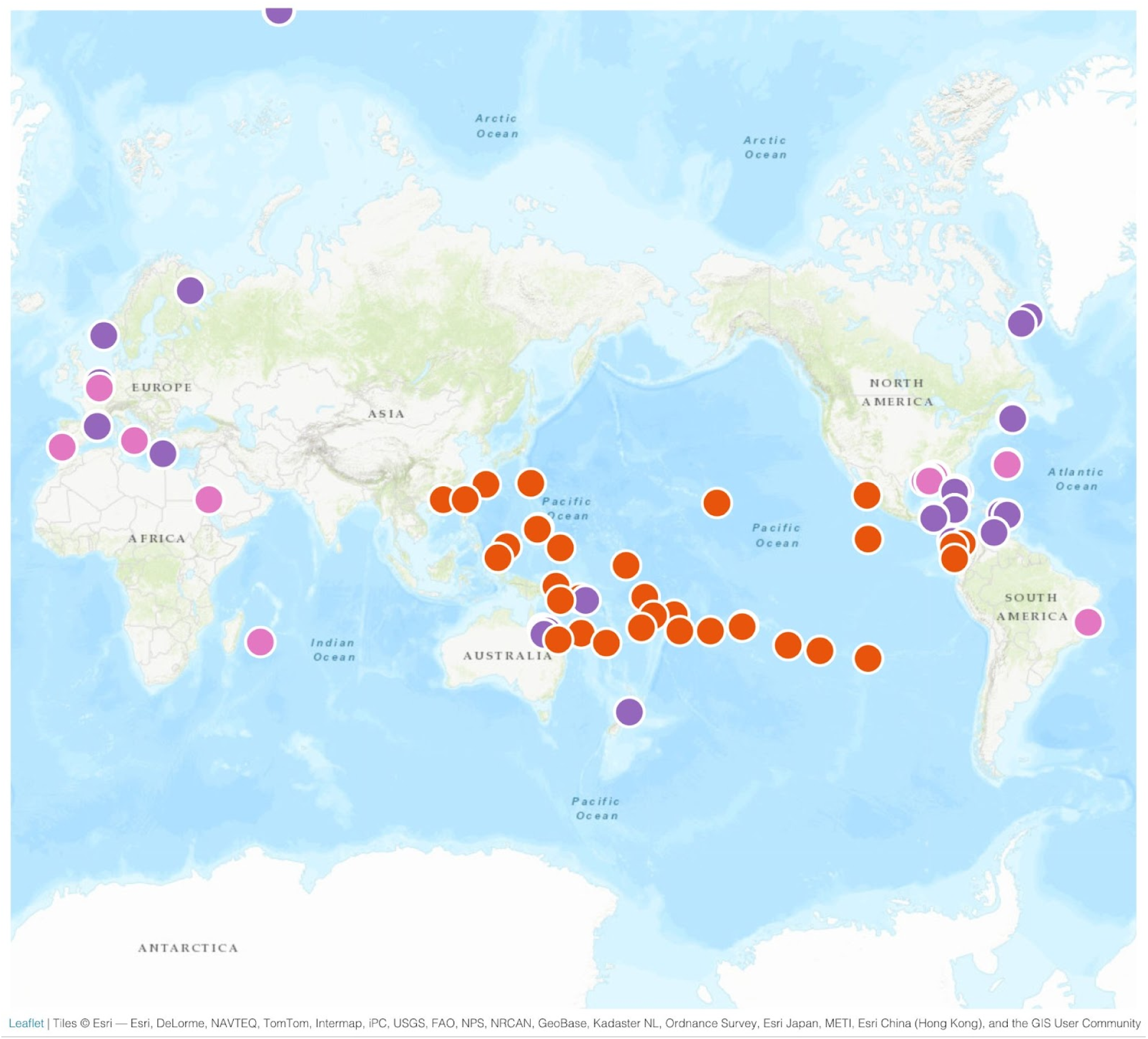
The coral reef metagenomes used in this study are distributed across the globe. We generated a genomic resource with global representation, by integrating the 820 coral metagenomes sampled by *Tara* Pacific (orange) with publicly available coral metagenomes (412) from 15 studies (pink) and sponge metagenomes (371) from 16 studies (purple).

**Supplementary Figure 2:**
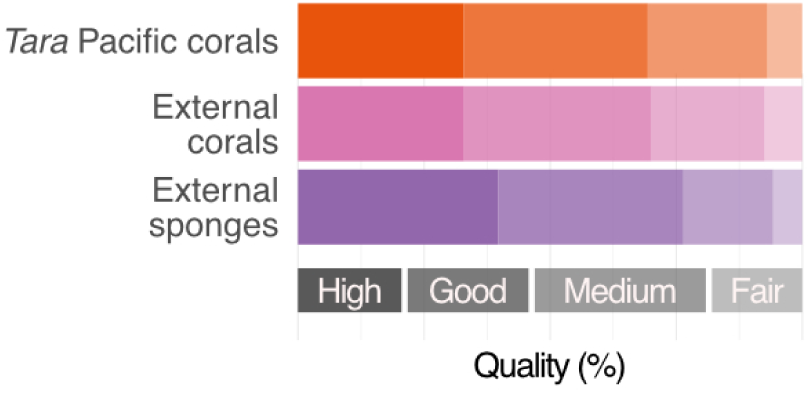
The reconstructed genomes display a high degree of completeness and little contamination. We attributed each metagenome-assembled genome (MAG) a quality score based on its completeness and degree of contamination (Methods). For each dataset, at least one third of all MAGs were of high quality (completeness ≥ 90% and contamination ≤ 5%) and more than two thirds were of good quality or higher (completeness ≥ 70% and contamination ≤ 10%), which compares favourably to previous large-scale efforts focusing on other microbiomes^26,28^.

**Supplementary Figure 3:**
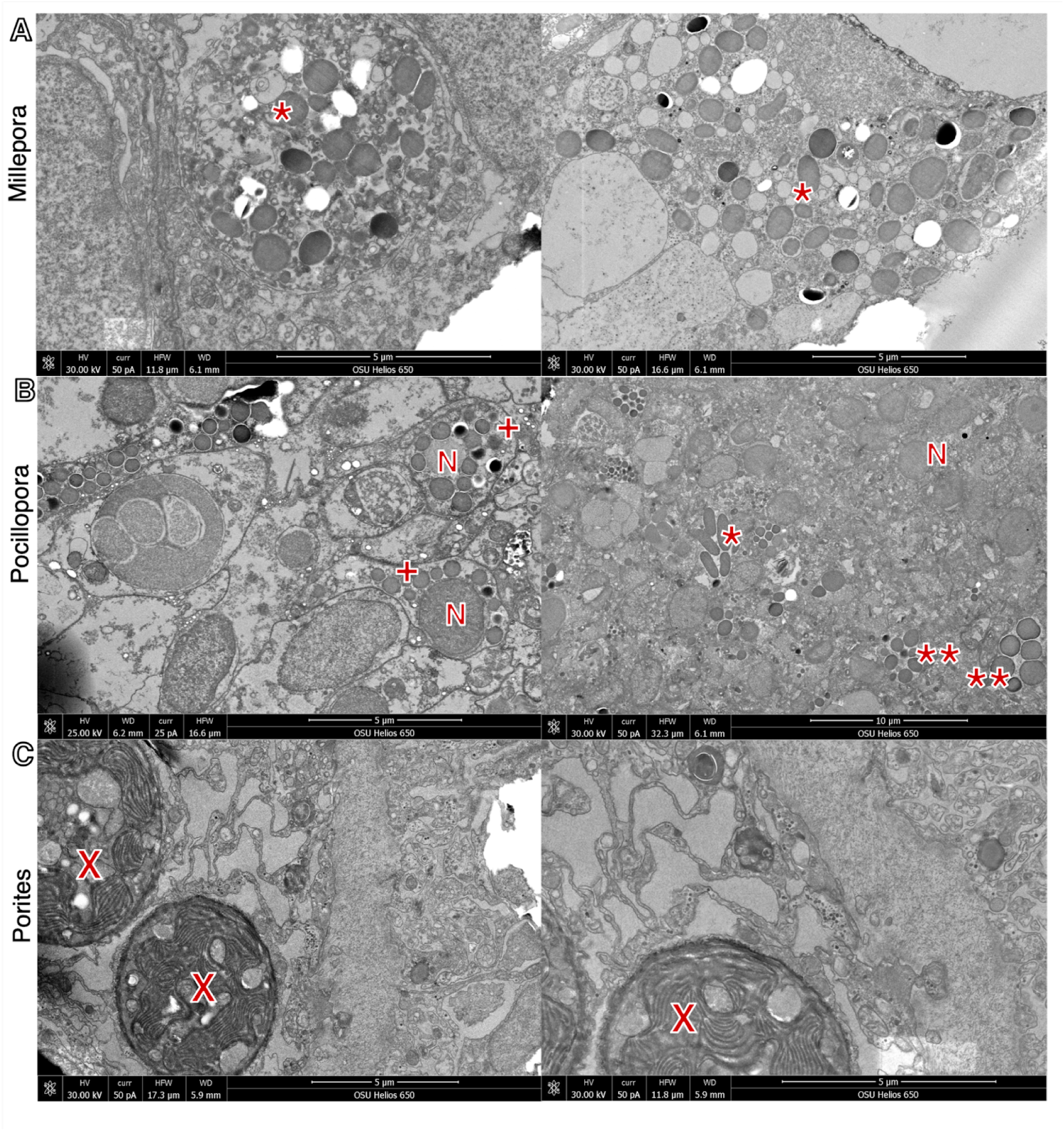
Transmission electron microscopy images show a high load of extracellular microorganisms in *Millepora.* By comparing the transmission electron microscopy images of the three coral genera (A) *Millepora*, (B) *Pocillopora*, and (C) *Porites* (sampled in Guam in 2016), we find differences in the number of microbial cells present as well as in the niche the microorganisms colonise. Annotations denote possible: (*) extracellular bacteria/archaea, (**) CAMAs (Coral-Associated Microbial Aggregates), (+) intracellular bacteria/archaea, (X) Symbiodinaceae (symbiont, including intracellular thylakoids), (N) coral nuclei.

**Supplementary Figure 4:**
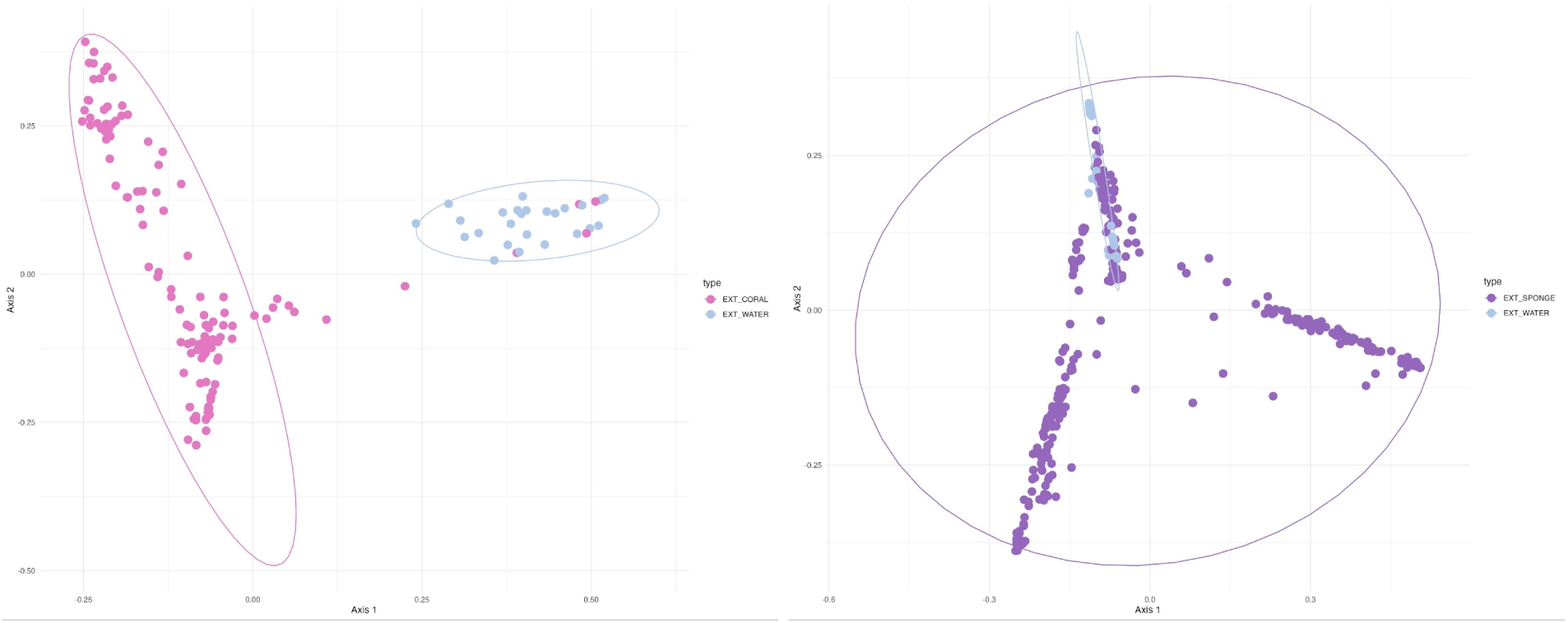
Corals and sponges host microbial communities that are distinct from those in the seawater. (left) A clear separation between the microbiomes of corals and seawater is found based on a Jaccard distance-based Principal Coordinate Analysis of the microbial species detected in coral and seawater metagenomes from publicly available coral studies (PERMANOVA, p-value≤0.001, R^2^=0.17, see Methods). **(right)** Similarly, the sponge metagenomes harbour microbial species that are distinct from those found in seawater metagenomes, although the difference is weaker than what was observed between coral and seawater metagenomes (PERMANOVA, p-value≤0.001, R^2^=0.08).

**Supplementary Figure 5:**
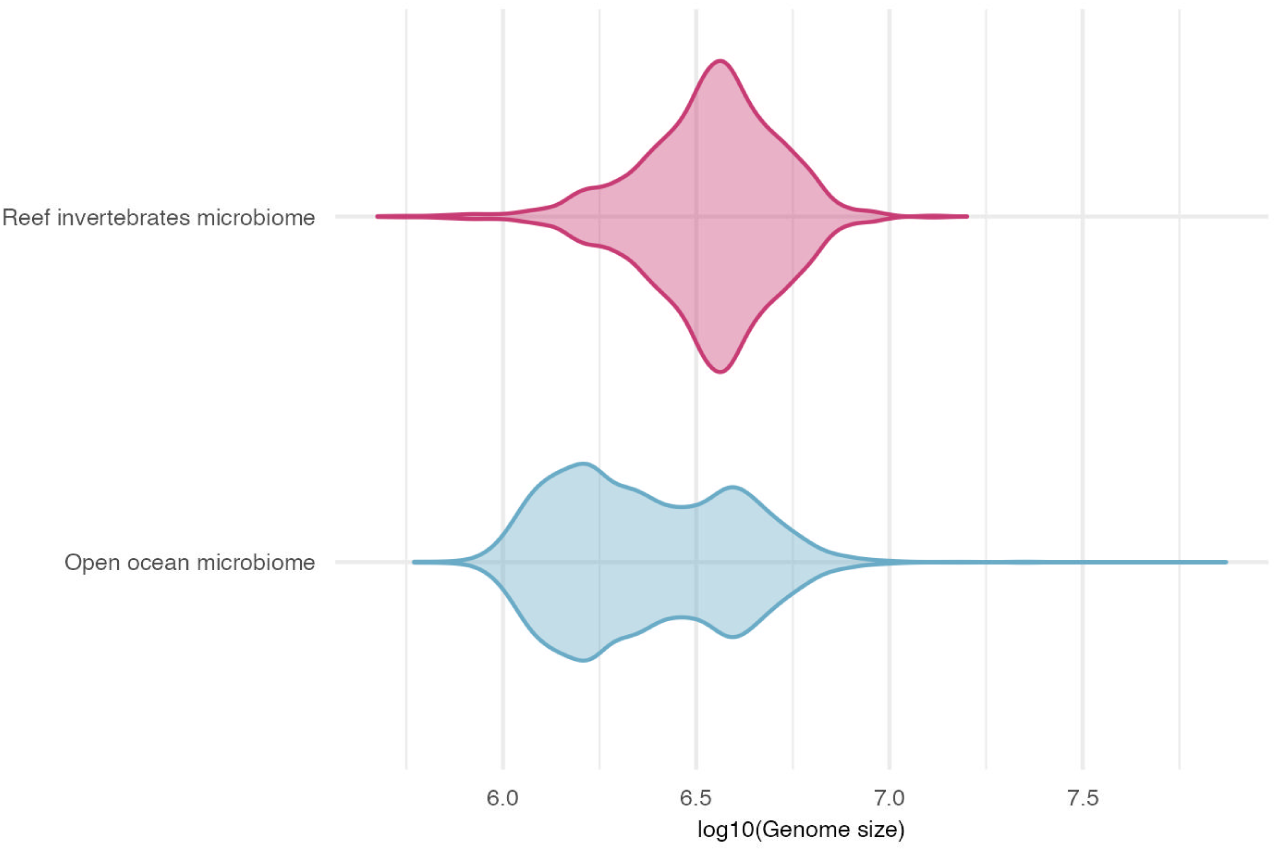
The estimated genome size of microbial species detected in the reef microbiome is larger than that of the ocean microbiome. After correcting the genome size estimates^28^ of any microbial species represented by at least one genome of more than 70% completeness, we found the median genome size of microbial species associated with reef invertebrates to be 3.6 Mbp, which was larger than that of the open ocean (2.2 Mbp; Wilcoxon-test p-value<2.2e-16).

**Supplementary Figure 6:**
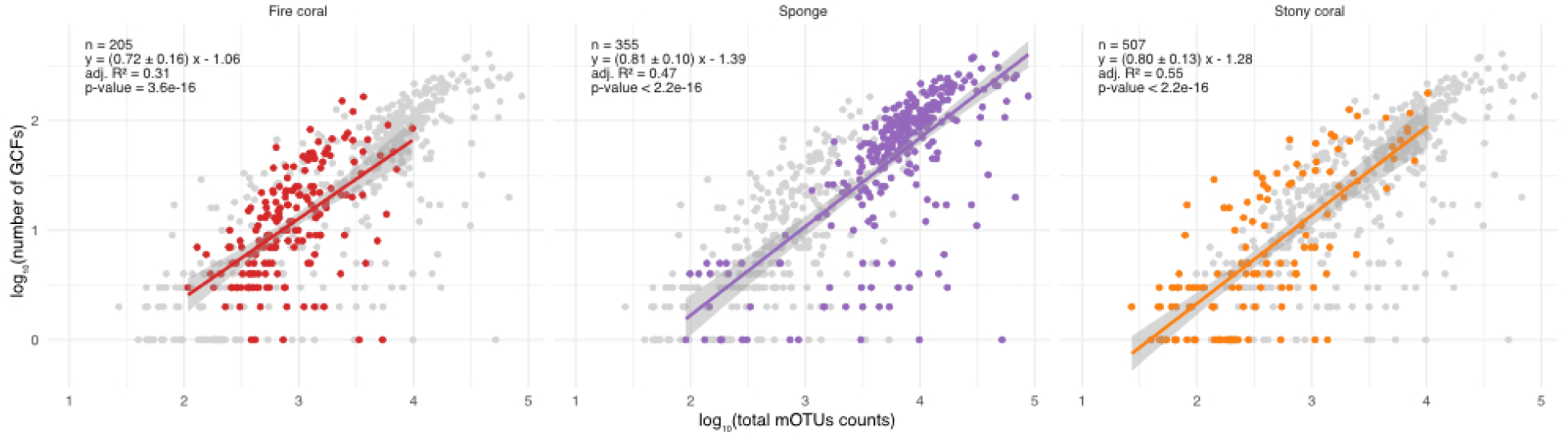
For all host groups, the number of GCFs depends on the sequencing depth. We sought to investigate the impact of the sequencing depth (as captured by the total mOTUs count, a proxy for the number of bacterial and archeal cells sequenced) on the number of GCFs identified in a metagenome across host groups. The higher number of GCFs identified in sponge metagenomes can be attributed to a higher number of microbial cells sequenced. The overall trends are similar across host groups after normalising by the sequencing effort (the 95% confidence intervals of the slope estimates overlap).

**Supplementary Figure 7:**
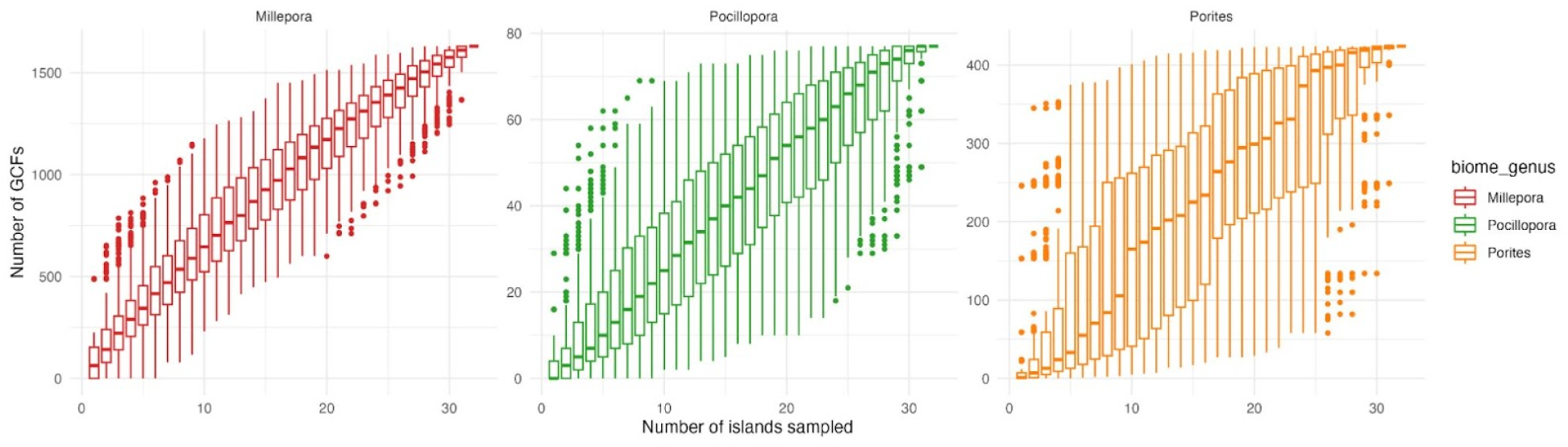
Increasing the geographic coverage would unveil additional GCFs. The rarefaction curves of the number of gene-cluster families (GCFs) identified across coral genera sampled by *Tara* Pacific (*Millepora*, *Pocillopora*, *Porites*) against the number of islands sampled (based on 500 randomisations) do not reach saturation and therefore suggest that we can uncover additional GCFs by further increasing the geographic range of sampling. The higher variation in *Porites* may indicate that the biosynthetic potential of its microbiome is more geographically heterogeneous.

**Supplementary Figure 8:**
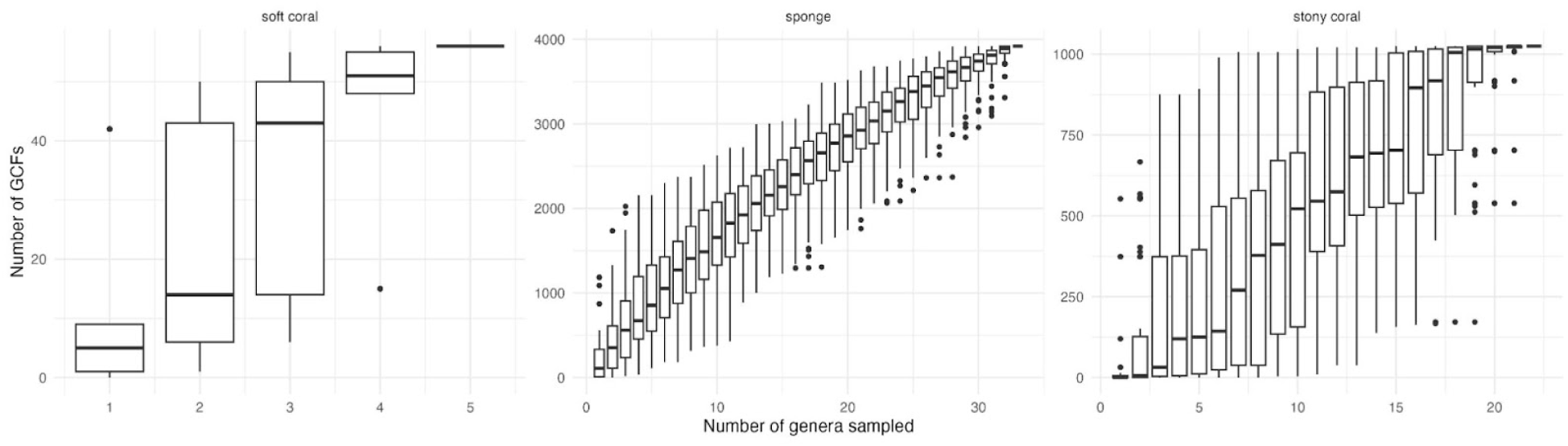
Sampling additional host species would unveil additional GCFs. The rarefaction curves of the number of gene-cluster families (GCFs) identified against the number of coral/sponge genera sampled by previous studies (based on 100 randomisations) do not reach saturation and therefore suggest that we can expect to uncover additional GCFs by increasing the host diversity range. The higher variation across soft and stony corals may suggest that the biosynthetic potential of their microbiome is more heterogeneous between genera. However, this higher variation may also result from the variable sampling effort and the various sampling protocols used (as they originate from many different studies) across genera.

**Supplementary Figure 9:**
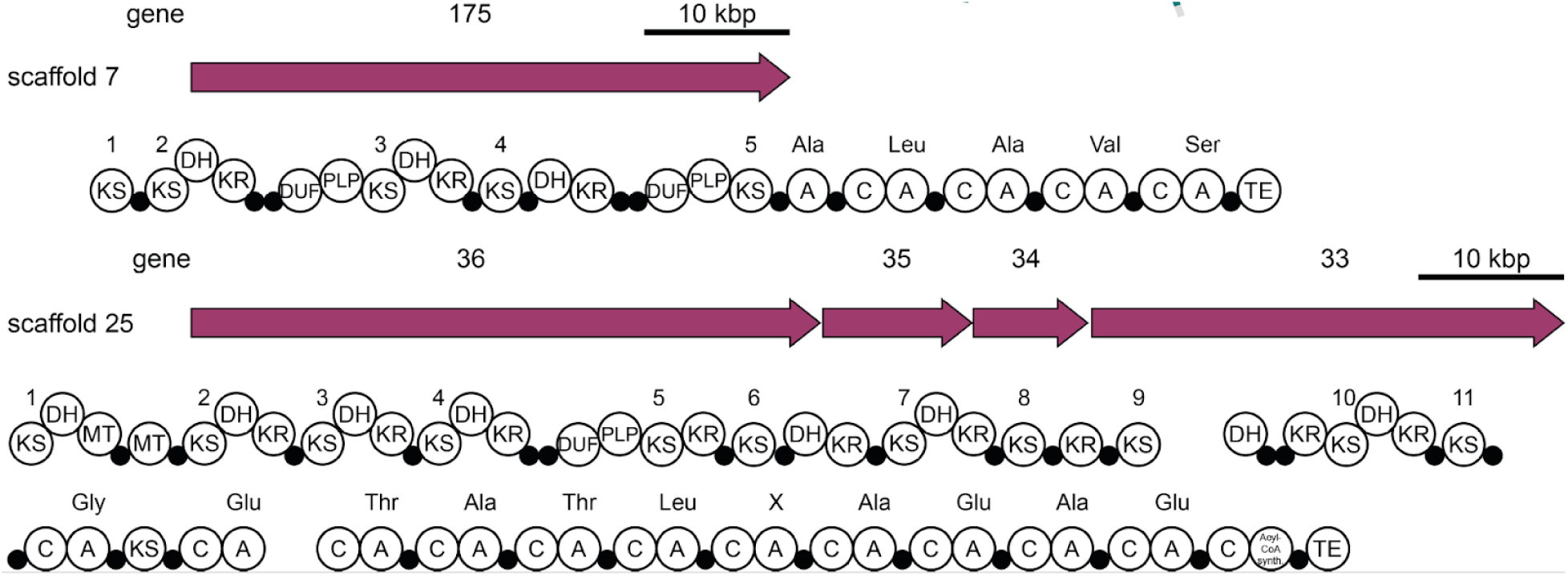
Complex *trans*-AT PKS clusters identified in an uncharacterised Acanthopleuribacteraceae species reconstructed from a fire coral (*Millepora*) metagenome. Two examples of architecturally unusual *trans*-AT PKS-NRPS BGCs from the *Millepora*-associated Acanthopleuribacteraceae MAG. The assembly lines comprise 10 and 24 modules, respectively, with a repeated occurrence of domains of unknown function (DUF) adjacent to pyridoxal phosphate-(PLP)dependent domains. Adapted from Leopod-Messer, Chepkirui et al., 2023^31^. All predicted BGCs for genomes from the two Acanthopleuribacteraceae spp. are listed in Supplementary Table 6.

## References

1. Smith, S. V. Coral-reef area and the contributions of reefs to processes and resources of the world’s oceans. Nature 273, 225–226 (1978).

2. Fisher, R. et al. Species richness on coral reefs and the pursuit of convergent global estimates. Curr. Biol. 25, 500–505 (2015).

3. Rocha, J., Peixe, L., Gomes, N. C. M. & Calado, R. Cnidarians as a source of new marine bioactive compounds—an overview of the last decade and future steps for bioprospecting. Mar. Drugs 9, 1860–1886 (2011).

4. Eddy, T. D. et al. Global decline in capacity of coral reefs to provide ecosystem services. One Earth 4, 1278–1285 (2021).

5. Knowlton, N. The future of coral reefs. Proc. Natl. Acad. Sci. USA 98, 5419–5425 (2001).

6. IPCC. Global Warming of 1.5°C. An IPCC Special Report on the impacts of global warming of 1.5°C above pre-industrial levels and related global greenhouse gas emission pathways, in the context of strengthening the global response to the threat of climate change, sustainable development, and efforts to eradicate poverty. (Cambridge University Press, 2018).

7. Galand, P. E. et al. Diversity of the Pacific Ocean coral reef microbiome. Nat. Commun. 14, 3039 (2023).

8. Muscatine, L. & Porter, J. W. Reef corals: Mutualistic symbioses adapted to nutrient-poor environments. BioScience 27, 454–460 (1977).

9. Lesser, M. P., Mazel, C. H., Gorbunov, M. Y. & Falkowski, P. G. Discovery of symbiotic nitrogen-fixing cyanobacteria in corals. Science 305, 997–1000 (2004).

10. Wiedenmann, J. et al. Reef-building corals farm and feed on their photosynthetic symbionts. Nature 620, 1018–1024 (2023).

11. Voolstra, C. R. & Ziegler, M. Adapting with microbial help: Microbiome flexibility facilitates rapid responses to environmental change. BioEssays 42, e2000004 (2020).

12. Ritchie, K. B. Regulation of microbial populations by coral surface mucus and mucus-associated bacteria. Mar. Ecol. Prog. Ser. 322, 1–14 (2006).

13. Vega Thurber, R., et al. Deciphering coral disease dynamics: Integrating host, microbiome, and the changing environment. Front. Ecol. Evol. 8, 575927 (2020).

14. Modolon, F., Barno, A. R., Villela, H. D. M. & Peixoto, R. S. Ecological and biotechnological importance of secondary metabolites produced by coral-associated bacteria. J. Appl. Microbiol. 129, 1441–1457 (2020).

15. Kobayashi, M. et al. Marine natural products. XXXIV. Trisindoline, a new antibiotic indole trimer, produced by a bacterium of Vibrio sp. separated from the marine sponge Hyrtios altum. Chem. Pharm. Bull. 42, 2449–2451 (1994).

16. Bertin, M. J. et al. Spongosine production by a *Vibrio harveyi* strain associated with the sponge *Tectitethya crypta*. J. Nat. Prod. 78, 493–499 (2015).

17. Romero, F. et al. Thiocoraline, a new depsipeptide with antitumor activity produced by a marine *Micromonospora*. J. Antibiot. 50, 734–737 (1997).

18. Newman, D. J. & Cragg, G. M. Marine natural products and related compounds in clinical and advanced preclinical trials. J. Nat. Prod. 67, 1216–1238 (2004).

19. Blockley, A., Elliott, D. R., Roberts, A. P. & Sweet, M. Symbiotic microbes from marine invertebrates: Driving a new era of natural product drug discovery. Diversity 9, 49 (2017).

20. Newman, D. J. From large-scale collections to the potential use of genomic techniques for supply of drug candidates. Front. Mar. Sci. 5, 401 (2018).

21. Sweet, M. et al. Insights into the cultured bacterial fraction of corals. mSystems 6, e0124920 (2021).

22. Pye, C. R., Bertin, M. J., Lokey, R. S., Gerwick, W. H. & Linington, R. G. Retrospective analysis of natural products provides insights for future discovery trends. Proc. Natl. Acad. Sci. USA 114, 5601–5606 (2017).

23. Blin, K. et al. antiSMASH 5.0: updates to the secondary metabolite genome mining pipeline. Nucleic Acids Res. 47, W81–W87 (2019).

24. Robinson, S. L., Piel, J. & Sunagawa, S. A roadmap for metagenomic enzyme discovery. Nat. Prod. Rep. 38, 1994–2023 (2021).

25. Pachiadaki, M. G. et al. Charting the complexity of the marine microbiome through single-cell genomics. Cell 179, 1623–1635 (2019).

26. Nayfach, S. et al. A genomic catalog of Earth’s microbiomes. Nat. Biotechnol. 39, 499–509 (2021).

27. Loureiro, C. et al. Comparative metagenomic analysis of biosynthetic diversity across sponge microbiomes highlights metabolic novelty, conservation, and diversification. mSystems 7, e00357–22 (2022).

28. Paoli, L. et al. Biosynthetic potential of the global ocean microbiome. Nature 607, 111–118 (2022).

29. van de Water, J. A., Tignat-Perrier, R., Allemand, D. & Ferrier-Pagès, C. Coral holobionts and biotechnology: from Blue Economy to coral reef conservation. Curr. Opin. Biotechnol. 74, 110–121 (2022).

30. 30. Planes, S., et al. The *Tara* Pacific expedition—A pan-ecosystemic approach of the ‘-omics’ complexity of coral reef holobionts across the Pacific Ocean. PLoS Biol. 17, e3000483 (2019).

31. Leopold-Messer, S. et al. Animal-associated marine Acidobacteria with a rich natural product repertoire. Chem.

32. Voolstra, C. R. et al. Disparate genetic divergence patterns in three corals across a pan-Pacific environmental gradient highlight species-specific adaptation. npj Biodiversity 2, 15 (2023).

33. Planes, S. & Allemand, D. Insights and achievements from the Tara Pacific expedition. Nat. Commun. 14, 3131 (2023).

34. Chaumeil, P.-A., Mussig, A. J., Hugenholtz, P. & Parks, D. H. GTDB-Tk: a toolkit to classify genomes with the Genome Taxonomy Database. Bioinformatics 36, 1925–1927 (2019).

35. Jain, C., Rodriguez-R, L. M., Phillippy, A. M., Konstantinidis, K. T. & Aluru, S. High throughput ANI analysis of 90K prokaryotic genomes reveals clear species boundaries. Nat. Commun. 9, 5114 (2018).

36. Ruscheweyh, H.-J. et al. Cultivation-independent genomes greatly expand taxonomic-profiling capabilities of mOTUs across various environments. Microbiome 10, 212 (2022).

37. Lombard, F. et al. Open science resources from the *Tara* Pacific expedition across coral reef and surface ocean ecosystems. Sci Data 10, 324 (2023).

38. Parks, D. H. et al. Recovery of nearly 8,000 metagenome-assembled genomes substantially expands the tree of life. Nat Microbiol 2, 1533–1542 (2017).

39. Garner, R. E. et al. A genome catalogue of lake bacterial diversity and its drivers at continental scale. Nat Microbiol 8, 1920–1934 (2023).

40. Rohwer, F., Seguritan, V., Azam, F. & Knowlton, N. Diversity and distribution of coral-associated bacteria. Mar. Ecol. Prog. Ser. 243, 1–10 (2002).

41. Sunagawa, S., Woodley, C. M. & Medina, M. Threatened corals provide underexplored microbial habitats. PLoS One 5, e9554 (2010).

42. Pollock, F. J. et al. Coral-associated bacteria demonstrate phylosymbiosis and cophylogeny. Nat. Commun. 9, 4921 (2018).

43. Sunagawa, S. et al. Structure and function of the global ocean microbiome. Science 348, 1261359 (2015).

44. Reshef, L., Koren, O., Loya, Y., Zilber-Rosenberg, I. & Rosenberg, E. The coral probiotic hypothesis. Environ. Microbiol. 8, 2068–2073 (2006).

45. Novichkov, P. S., Wolf, Y. I., Dubchak, I. & Koonin, E. V. Trends in prokaryotic evolution revealed by comparison of closely related bacterial and archaeal genomes. J. Bacteriol. 191, 65–73 (2009).

46. Giovannoni, S. J., Cameron Thrash, J. & Temperton, B. Implications of streamlining theory for microbial ecology. ISME J. 8, 1553–1565 (2014).

47. Kautsar, S. A., van der Hooft, J. J. J., de Ridder, D. & Medema, M. H. BiG-SLiCE: A highly scalable tool maps the diversity of 1.2 million biosynthetic gene clusters. GigaScience 10, giaa154 (2021).

48. Ghurye, J. S., Cepeda-Espinoza, V. & Pop, M. Metagenomic assembly: Overview, challenges and applications. Yale J. Biol. Med. 89, 353–362 (2016).

49. Kautsar, S. A. et al. MIBiG 2.0: a repository for biosynthetic gene clusters of known function. Nucleic Acids Res. 48, D454–D458 (2020).

50. Kautsar, S. A., Blin, K., Shaw, S., Weber, T. & Medema, M. H. BiG-FAM: the biosynthetic gene cluster families database. Nucleic Acids Res. 49, D490–D497 (2021).

51. Almeida, J. F. et al. Marine sponge and octocoral-associated bacteria show versatile secondary metabolite biosynthesis potential and antimicrobial activities against human pathogens. Mar. Drugs 21, 34 (2022).

52. Carroll, A. R., Copp, B. R., Davis, R. A., Keyzers, R. A. & Prinsep, M. R. Marine natural products. Nat. Prod. Rep. 39, 1122–1171 (2022).

53. Schwarzer, D., Finking, R. & Marahiel, M. A. Nonribosomal peptides: from genes to products. Nat. Prod. Rep. 20, 275–287 (2003).

54. Dubé, C. E. et al. Naturally occurring fire coral clones demonstrate a genetic and environmental basis of microbiome composition. Nat. Commun. 12, 6402 (2021).

55. Crits-Christoph, A., Diamond, S., Butterfield, C. N., Thomas, B. C. & Banfield, J. F. Novel soil bacteria possess diverse genes for secondary metabolite biosynthesis. Nature 558, 440–444 (2018).

56. Hemmerling, F. & Piel, J. Strategies to access biosynthetic novelty in bacterial genomes for drug discovery. Nat. Rev. Drug Discov. 21, 359–378 (2022).

57. Freeman, M. F., Vagstad, A. L. & Piel, J. Polytheonamide biosynthesis showcasing the metabolic potential of sponge-associated uncultivated ‘Entotheonella’ bacteria. Curr. Opin. Chem. Biol. 31, 8–14 (2016).

58. 58. Kogawa, M., et al. Single-cell metabolite detection and genomics reveals uncultivated talented producer. *PNAS Nexus* 1, gab007 (2022).

59. Helfrich, E. J. N. & Piel, J. Biosynthesis of polyketides by trans-AT polyketide synthases. Nat. Prod. Rep. 33, 231–316 (2016).

60. Hoeksema, 44B. W. & Cairns, S. Word list of Scleractinia, Accessed on 2023-11-20 at. https://www.marinespecies.org/scleractinia.

61. Pesant, S., et al. Tara Pacific samples provenance and environmental context - version 2. (2020) doi:10.5281/ZENODO.4068292.

62. Belser, C. et al. Integrative omics framework for characterization of coral reef ecosystems from the *Tara Pacific* expedition. Sci Data 10, 326 (2023).

63. Pfeifer, C. R. et al. Quantitative analysis of mouse pancreatic islet architecture by serial block-face SEM. J. Struct. Biol. 189, 44–52 (2015).

64. Leblud, J., Moulin, L., Batigny, A., Dubois, P. & Grosjean, P. Artificial coral reef mesocosms for ocean acidification investigations. Biogeosci. Discuss. 11, 15463–15505 (2014).

65. Meyer, J. L., Gunasekera, S. P., Scott, R. M., Paul, V. J. & Teplitski, M. Microbiome shifts and the inhibition of quorum sensing by Black Band Disease cyanobacteria. ISME J. 10, 1204–1216 (2016).

66. Cai, L. et al. Metagenomic analysis reveals a green sulfur bacterium as a potential coral symbiont. Sci. Rep. 7, 9320 (2017).

67. Sato, Y. et al. Unraveling the microbial processes of black band disease in corals through integrated genomics. Sci. Rep. 7, 40455 (2017).

68. Wang, L. et al. Corals and their microbiomes are differentially affected by exposure to elevated nutrients and a natural thermal anomaly. Front. Mar. Sci. 5, 101 (2018).

69. Robbins, S. J. et al. A genomic view of the reef-building coral *Porites lutea* and its microbial symbionts. Nat. Microbiol. 4, 2090–2100 (2019).

70. Yang, S.-H. et al. Metagenomic, phylogenetic, and functional characterization of predominant endolithic green sulfur bacteria in the coral *Isopora palifera*. Microbiome 7, 3 (2019).

71. Lima, L. F. O. et al. Modeling of the coral microbiome: the influence of temperature and microbial network. mBio 11, e02691–19 (2020).

72. Messyasz, A. et al. Coral bleaching phenotypes associated with differential abundances of nucleocytoplasmic large DNA viruses. Front. Mar. Sci. 7, 555474 (2020).

73. Ngugi, D. K., Ziegler, M., Duarte, C. M. & Voolstra, C. Genomic blueprint of glycine betaine metabolism in coral metaorganisms and their contribution to reef nitrogen budgets. iScience 23, 101120 (2020).

74. Vohsen, S. A. et al. Deep-sea corals provide new insight into the ecology, evolution, and the role of plastids in widespread apicomplexan symbionts of anthozoans. Microbiome 8, 34 (2020).

75. Santoro, E. P. et al. Coral microbiome manipulation elicits metabolic and genetic restructuring to mitigate heat stress and evade mortality. Sci Adv 7, eabg3088 (2021).

76. Keller-Costa, T. et al. Metagenomic insights into the taxonomy, function, and dysbiosis of prokaryotic communities in octocorals. Microbiome 9, 72 (2021).

77. Cárdenas, A. et al. Greater functional diversity and redundancy of coral endolithic microbiomes align with lower coral bleaching susceptibility. ISME J. 16, 2406–2420 (2022).

78. Palladino, G. et al. Metagenomic shifts in mucus, tissue and skeleton of the coral *Balanophyllia europaea* living along a natural CO_2_ gradient. ISME Commun. 2, 65 (2022).

79. Agarwal, V. et al. Metagenomic discovery of polybrominated diphenyl ether biosynthesis by marine sponges. Nat. Chem. Biol. 13, 537–543 (2017).

80. Jahn, M. T. et al. A phage protein aids bacterial symbionts in eukaryote immune evasion. Cell Host Microbe 26, 542–550 (2019).

81. Busch, K. et al. Microbial diversity of the glass sponge *Vazella pourtalesii* in response to anthropogenic activities. Conserv. Genet. 21, 1001–1010 (2020).

82. Engelberts, J. P. et al. Characterization of a sponge microbiome using an integrative genome-centric approach. ISME J. 14, 1100–1110 (2020).

83. Glasl, B. et al. Comparative genome-centric analysis reveals seasonal variation in the function of coral reef microbiomes. ISME J. 14, 1435–1450 (2020).

84. Pascelli, C. et al. Viral ecogenomics across the Porifera. Microbiome 8, 144 (2020).

85. Storey, M. A. et al. Metagenomic exploration of the marine sponge *Mycale hentscheli* uncovers multiple polyketide-producing bacterial symbionts. mBio 11, e02997–19 (2020).

86. Nguyen, N. A. et al. An obligate peptidyl brominase underlies the discovery of highly distributed biosynthetic gene clusters in marine sponge microbiomes. J. Am. Chem. Soc. 143, 10221–10231 (2021).

87. Rusanova, A., Fedorchuk, V., Toshchakov, S., Dubiley, S. & Sutormin, D. An interplay between viruses and bacteria associated with the white sea sponges revealed by metagenomics. Life 12, 25 (2021).

88. Robbins, S. J. et al. A genomic view of the microbiome of coral reef demosponges. ISME J. 15, 1641–1654 (2021).

89. Dharamshi, J. E. et al. Genomic diversity and biosynthetic capabilities of sponge-associated chlamydiae. ISME J. 16, 2725–2740 (2022).

90. Kelly, J. B., Carlson, D. E., Low, J. S. & Thacker, R. W. Novel trends of genome evolution in highly complex tropical sponge microbiomes. Microbiome 10, 164 (2022).

91. Morganti, T. M. et al. Giant sponge grounds of Central Arctic seamounts are associated with extinct seep life. Nat. Commun. 13, 638 (2022).

92. Pankey, M. S. et al. Cophylogeny and convergence shape holobiont evolution in sponge–microbe symbioses. *Nat*. Ecol. Evol. 6, 750–762 (2022).

93. Engelberts, J. P. et al. Metabolic reconstruction of the near complete microbiome of the model sponge *Ianthella basta*. Environ. Microbiol. 25, 646–660 (2023).

94. Thompson, C. C. et al. Genomic taxonomy of vibrios. BMC Evol. Biol. 9, 258 (2009).

95. Kimes, N. E. et al. Temperature regulation of virulence factors in the pathogen *Vibrio coralliilyticus*. ISME J. 6, 835–846 (2012).

96. Bondarev, V. et al. The genus *Pseudovibrio* contains metabolically versatile bacteria adapted for symbiosis. Environ. Microbiol. 15, 2095–2113 (2013).

97. Ushijima, B., et al. *Vibrio coralliilyticus* strain OCN008 is an etiological agent of acute *Montipora* white syndrome. Appl. Environ. Microbiol. 80, 2102–2109 (2014).

98. Asahina, A. Y. & Hadfield, M. G. Draft genome sequence of *Pseudoalteromonas luteoviolacea* HI1, determined using Roche 454 and PacBio single-molecule real-time hybrid sequencing. Genome Announc. 3, e01590–14 (2015).

99. Meyer, J. L. et al. Draft genome sequence of *Halomonas meridiana* R1t3 isolated from the surface microbiota of the Caribbean Elkhorn coral *Acropora palmata*. Stand. Genomic Sci. 10, 75 (2015).

100. Ding, J.-Y., Shiu, J.-H., Chen, W.-M., Chiang, Y.-R. & Tang, S.-L. Genomic insight into the host-endosymbiont relationship of *Endozoicomonas montiporae* CL-33^T^ with its coral host. Front. Microbiol. 7, 251 (2016).

101. Franco, T., Califano, G., Gonçalves, A. C., Cúcio, C. & Costa, R. Draft genome sequence of *Vibrio* sp. strain Evh12, a bacterium retrieved from the gorgonian coral *Eunicella verrucosa*. Genome Announc. 4, e01729–15 (2016).

102. Keller-Costa, T., Silva, R., Lago-Lestón, A. & Costa, R. Genomic insights into *Aquimarina* sp. strain EL33, a bacterial symbiont of the gorgonian coral *Eunicella labiata*. Genome Announc. 4, e00855–16 (2016).

103. Wan, X., Miller, J. M., Rowley, S. J., Hou, S. & Donachie, S. P. Draft genome sequence of a novel *Luteimonas* sp. strain from coral mucus, Hawai‘i. Genome Announc. 4, e01228–16 (2016).

104. Henao, J. et al. Genome sequencing of three bacteria associated to black band disease from a Colombian reef-building coral. Genom Data 11, 73–74 (2017).

105. Braun, D. R. et al. Complete genome sequence of *Dietzia* sp. strain WMMA184, a marine coral-associated bacterium. Genome Announc. 6, e01582–17 (2018).

106. Kumari, P., Badhai, J. & Das, S. K. Draft genome sequence of *Marinomonas fungiae* strain AN44^T^ (JCM 18476^T^), isolated from the coral *Fungia echinata* from the Andaman Sea. Genome Announc. 6, e00112–18 (2018).

107. Raimundo, I., Silva, S. G., Costa, R. & Keller-Costa, T. Bioactive secondary metabolites from octocoral-associated microbes: New chances for blue growth. Mar. Drugs 16, 485 (2018).

108. Rodrigues, G. N., Lago-Lestón, A., Costa, R. & Keller-Costa, T. Draft genome sequence of *Labrenzia* sp. strain EL143, a coral-associated Alphaproteobacterium with versatile symbiotic living capability and strong halogen degradation potential. Genome Announc. 6, e00132–18 (2018).

109. Silva, S. G., Lago-Lestón, A., Costa, R. & Keller-Costa, T. Draft genome sequence of *Sphingorhabdus* sp. strain EL138, a metabolically versatile Alphaproteobacterium isolated from the gorgonian coral *Eunicella labiata*. Genome Announc. 6, e00142–18 (2018).

110. Tandon, K., Chiang, P.-W., Chen, W.-M. & Tang, S.-L. Draft genome sequence of *Endozoicomonas acroporae* strain Acr-14^T^, isolated from *Acropora* coral. Genome Announc. 6, e01576–17 (2018).

111. Deb, S., Badhai, J. & Das, S. K. Draft genome sequences of two *Vibrio fortis* strains isolated from coral (*Fungia* sp.) from the Andaman Sea. Microbiol Resour Announc 9, e01225–19 (2020).

112. Li, J. et al. Cultured bacteria provide insight into the functional potential of the coral-associated microbiome. mSystems 7, e00327–22 (2022).

113. Shi, S.-B., Cui, L.-Q., Zeng, Q., Long, L.-J. & Tian, X.-P. *Nocardioides coralli* sp. nov., an actinobacterium isolated from stony coral in the South China Sea. Int. J. Syst. Evol. Microbiol. 72, 005342 (2022).

114. Nurk, S., Meleshko, D., Korobeynikov, A. & Pevzner, P. A. metaSPAdes: a new versatile metagenomic assembler. Genome Res. 27, 824–834 (2017).

115. Li, H. & Durbin, R. Fast and accurate short read alignment with Burrows-Wheeler transform. Bioinformatics 25, 1754–1760 (2009).

116. Kang, D. D. et al. MetaBAT 2: an adaptive binning algorithm for robust and efficient genome reconstruction from metagenome assemblies. PeerJ 7, e7359 (2019).

117. Parks, D. H., Imelfort, M., Skennerton, C. T., Hugenholtz, P. & Tyson, G. W. CheckM: assessing the quality of microbial genomes recovered from isolates, single cells, and metagenomes. Genome Res. 25, 1043–1055 (2015).

118. Eren, A. M. et al. Anvi’o: an advanced analysis and visualization platform for ’omics data. PeerJ 3, e1319 (2015).

119. Bowers, R. M. et al. Minimum information about a single amplified genome (MISAG) and a metagenome-assembled genome (MIMAG) of bacteria and archaea. Nat. Biotechnol. 35, 725–731 (2017).

120. Olm, M. R., Brown, C. T., Brooks, B. & Banfield, J. F. dRep: a tool for fast and accurate genomic comparisons that enables improved genome recovery from metagenomes through de-replication. ISME J. 11, 2864–2868 (2017).

121. Olm, M. R. et al. Consistent metagenome-derived metrics verify and delineate bacterial species boundaries. mSystems 5, e00731–19 (2020).

122. Parks, D. H. et al. A standardized bacterial taxonomy based on genome phylogeny substantially revises the tree of life. Nat. Biotechnol. 36, 996–1004 (2018).

123. Hyatt, D. et al. Prodigal: prokaryotic gene recognition and translation initiation site identification. BMC Bioinformatics 11, 119 (2010).

124. Laslett, D. & Canback, B. ARAGORN, a program to detect tRNA genes and tmRNA genes in nucleotide sequences. Nucleic Acids Res. 32, 11–16 (2004).

125. Milanese, A. et al. Microbial abundance, activity and population genomic profiling with mOTUs2. Nat. Commun. 10, 1014 (2019).

126. Paradis, E. & Schliep, K. ape 5.0: an environment for modern phylogenetics and evolutionary analyses in R. Bioinformatics 35, 526–528 (2019).

127. Salazar, G. et al. Gene expression changes and community turnover differentially shape the global ocean metatranscriptome. Cell 179, 1068–1083 (2019).

128. Fu, L., Niu, B., Zhu, Z., Wu, S. & Li, W. CD-HIT: accelerated for clustering the next-generation sequencing data. Bioinformatics 28, 3150–3152 (2012).

129. Huerta-Cepas, J. et al. Fast genome-wide functional annotation through orthology assignment by eggNOG-mapper. Mol. Biol. Evol. 34, 2115–2122 (2017).

130. Huerta-Cepas, J. et al. eggNOG 5.0: a hierarchical, functionally and phylogenetically annotated orthology resource based on 5090 organisms and 2502 viruses. Nucleic Acids Res. 47, D309–D314 (2019).

131. Kanehisa, M. & Goto, S. KEGG: Kyoto Encyclopedia of Genes and Genomes. Nucleic Acids Res. 28, 27–30 (2000).

132. Buchfink, B., Xie, C. & Huson, D. H. Fast and sensitive protein alignment using DIAMOND. Nat. Methods 12, 59–60 (2015).

133. Tatusova, T. et al. NCBI prokaryotic genome annotation pipeline. Nucleic Acids Res. 44, 6614–6624 (2016).

134. Gavriilidou, A. et al. Compendium of specialized metabolite biosynthetic diversity encoded in bacterial genomes. Nat Microbiol 7, 726–735 (2022).

135. Navarro-Muñoz, J. C. et al. A computational framework to explore large-scale biosynthetic diversity. Nat. Chem. Biol. 16, 60–68 (2020).

136. Ruscheweyh, H.-J., et al. Tara Pacific 16S rRNA data analysis release. (2022) doi:10.5281/ZENODO.4073268.

137. Conway, J. R., Lex, A. & Gehlenborg, N. UpSetR: an R package for the visualization of intersecting sets and their properties. Bioinformatics 33, 2938–2940 (2017).

138. Lex, A., Gehlenborg, N., Strobelt, H., Vuillemot, R. & Pfister, H. UpSet: Visualization of intersecting sets. IEEE Trans. Vis. Comput. Graph. 20, 1983–1992 (2014).

139. Krassowski, M. ComplexUpset. Preprint at 10.5281/zenodo.3700590 (2020).

140. Paoli, L., Wiederkehr, F., Ruscheweyh, H.-J. & Sunagawa, S. Genome-resolved diversity and biosynthetic potential of the coral reef microbiome. (2023) doi:10.5281/ZENODO.10182967.

